# Crumbs complex-directed apical membrane dynamics in epithelial cell ingression

**DOI:** 10.1101/2022.03.13.484184

**Authors:** Sérgio Simões, Gerald Lerchbaumer, Milena Pellikka, Paraskevi Giannatou, Thomas Lam, Dohyun Kim, Jessica Yu, David ter Stal, Kenana Al Kakouni, Rodrigo Fernandez-Gonzalez, Ulrich Tepass

## Abstract

Epithelial cells often leave their tissue context and ingress to form new cell types or acquire migratory ability to move to distant sites during development and tumor progression. Cell lose their apical membrane and epithelial adherens junctions during ingression. However, how factors that organize apical-basal polarity contribute to ingression is unknown. Here, we show that the dynamic regulation of the apical Crumbs polarity complex is crucial for normal neural stem cell ingression. Crumbs endocytosis and recycling allow ingression to occur in a normal timeframe. During early ingression, Crumbs and its complex partner the RhoGEF Cysts support myosin and apical constriction to ensure robust ingression dynamics. During late ingression, the E3-ubiquitin ligase Neuralized facilitates the disassembly of the Crumbs complex and the rapid endocytic removal of the apical cell domain. Our findings reveal a mechanism integrating cell fate, apical polarity, endocytosis, vesicle trafficking, and actomyosin contractility to promote cell ingression, a fundamental morphogenetic process observed in animal development and cancer.

## Introduction

The loss of apical-basal polarity and cell junctions are key early steps in epithelial to mesenchymal transitions (EMT) when cells leave the epithelium [Hay, 1995; Nieto et al., 2016; Campbell, 2018; Yang et al., 2020; Lambert and Weinberg, 2021]. EMTs drive cell escape during epithelial tumour progression and are frequent in animal development. Examples include the ingression of primary mesenchyme cells in the sea urchin embryo, the formation of mesoderm and endoderm during bird and mouse gastrulation, and the emergence of the neural crest in the vertebrate embryo [Shook and Keller, 2003; Lim and Thiery, 2012; Serrano-Najera and Weijer, 2020; Sheng, 2021]. In the Drosophila embryo, EMT is observed during mesoderm and endoderm development [Campbell et al., 2011; Gheisari et al., 2020] and in the neuroepithelium [Hartenstein and Wodarz, 2013]. We use the ingression of neural stem cells (or neuroblasts, NBs) as a model to study the mechanisms regulating an EMT-like process [Simoes et al., 2017; An et al., 2017]. Our analysis revealed that the reduction of the apical surface of NBs is driven by 10-12 oscillating ratcheted contractions of actomyosin and the progressive loss of adherens junctions (AJs) to neighboring neuroepithelial cells.

Epithelial polarity is governed by a network of polarity factors [Tepass, 2012; Rodriguez-Boulan and Macara, 2014; Pickett et al., 2019]. Here we investigate how the Crumbs (Crb) complex contributes to the loss of the apical domain – the apical membrane and AJs – during ingression. Crb is a transmembrane protein that governs apical membrane stability and the integrity of the circumferential AJs [Tepass et al., 1990; Wodarz et al., 1995; Tepass, 1996; Grawe et al., 1996; Silver et al., 2019]. Crb plays multiple roles in support of apical-basal polarity. It interacts with Moesin and the Spectrin cytoskeleton to support the apical cytocortex [Pellikka et al., 2002; Medina et al., 2002]. It interacts with the apical Par polarity complex including atypical Protein Kinase C (aPKC), which prevents apical enrichment of basolateral polarity proteins such as Lethal giant larvae or Yurt [Tepass, 2012; Morais-de-Sa et al., 2010, Pichaud et al., 2019; Gamblin et al., 2014]. Crb and its binding partners also recruit the RhoGEF Cysts (Cyst) to the apical junction. Cyst supports Rho1 activity and junctional myosin II, thus coupling Crb to junctional myosin stability [Silver et al., 2019]. Here, we have investigated how Crb contributes to the ordered reduction and ultimate removal of the apical domain during NB ingression.

## Results

### Crumbs and E-cadherin are lost from the NB apical domain with different kinetics

Neurogenesis in the Drosophila embryo is initiated by the emergence of NBs from the neuroectodermal epithelium (Fig. 1A) [Hartenstein and Wodarz, 2013]. Most NBs ingress from the epithelium as individual cells, a ∼30 minute process during which NBs lose apical-basal polarity [Simoes et al., 2017; An et al., 2017]. NBs lose their apical membrane including Crb as they ingress [Tepass et al., 1990], and remove their apical E-cadherin (Ecad)-based AJ [Tepass et al., 1996]. (Fig. 1B). Interestingly, we noticed a marked difference in how these two transmembrane proteins are lost. The concentration of junctional Ecad remained relatively constant during ingression showing only an 1.12+/-0.28 fold change over time (Fig. 1B,C) [Simoes et al., 2017], whereas junctional Crb increased 1.35+/-0.65 fold (Fig. 1B,C). In addition, while total apical Ecad levels declined in conjunction with apical area reduction, Crb total apical levels remained constant during the first 20 minutes (= ‘early ingression’) before declining rapidly during the last 10 minutes of apical area loss (= ‘late ingression’) (Fig. 1A-D). This suggests that Ecad and Crb are removed from the apical domain of NBs by distinct mechanisms.

**Figure 1.**
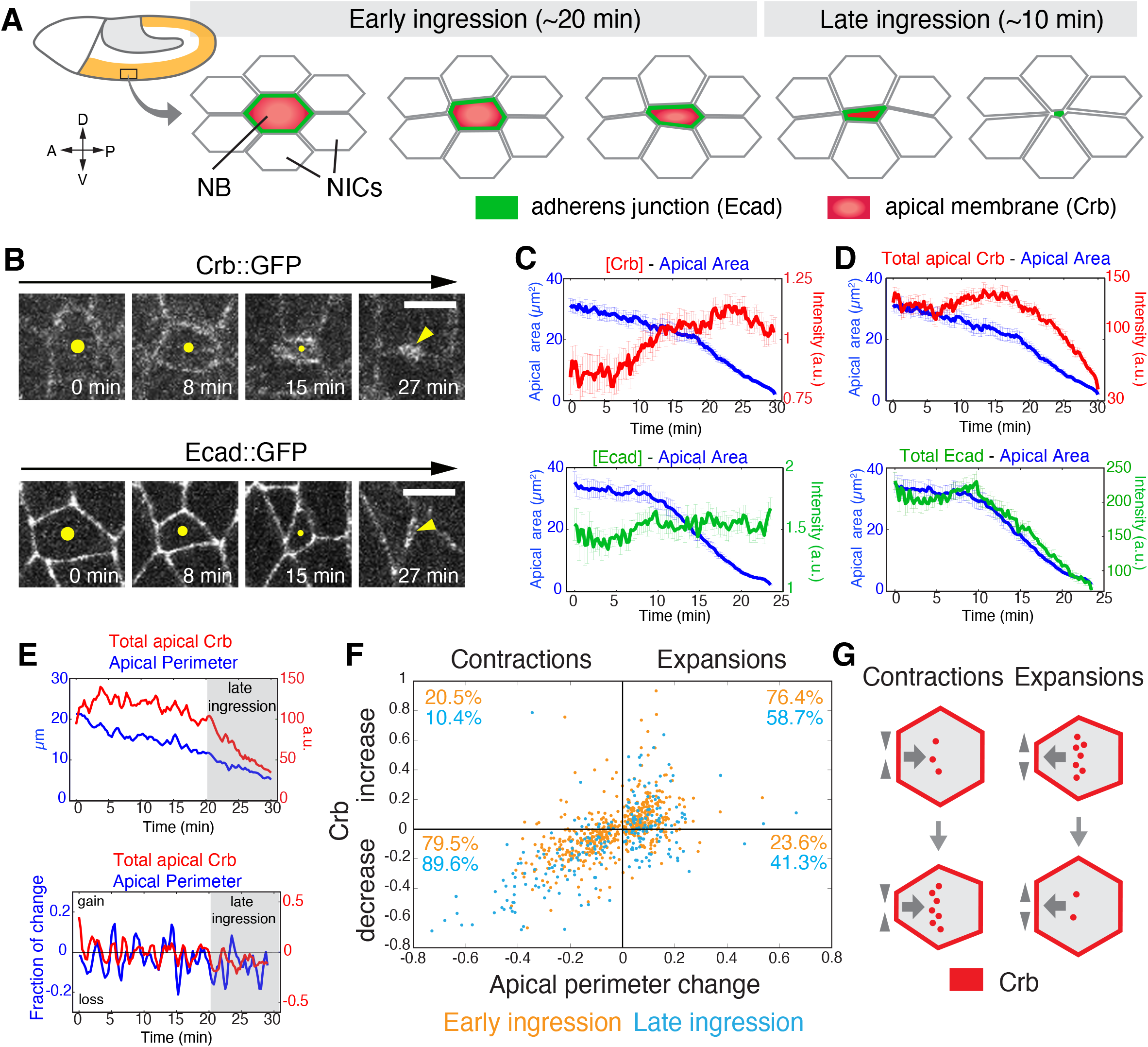
Crb persists during early NB ingression and undergoes rapid removal during late ingression. **(A)** Schematic of gastrulating Drosophila embryo (stage 8) when S1 NBs ingress. NICs, non-ingressing cells. **(B)** Apical surface of ingressing live NBs (yellow dots or arrowheads) expressing endo-Crb::GFP or endo-Ecad::GFP. Time after onset of ingression. Scale bars, 5 µm. **(C-D)** Junctional Crb::GFP and Ecad::GFP fluorescence (average pixel intensity at cell boundaries; C) and total apical protein levels (fluorescence intensity per cell; D). Blue lines represent average apical area. 58 NBs, 12 embryos for endo-Crb::GFP and 30 NBs, 6 embryos for endo-Ecad::GFP. T=0 min, the onset of ingression. Error bars are SEM. **(E)** Apical perimeter and total apical Crb::GFP during ingression (upper panel; T=0 min, onset of ingression) and fraction of change in apical perimeter and total Crb::GFP during ingression of a representative NB. Apical Crb levels decrease (rate <0) during apical contractions and increase (rate >0) during apical expansions. Apical Crb losses overtake gains during late ingression. **(F)** Scatter plot showing the relationship between the fraction of perimeter change during apical contractions or expansions and the corresponding fraction of total apical Crb change (relative Crb increase or decrease). N values as in Fig. S1A. **(G)** Schematic interpretation of data in (F) illustrating preferential Crb membrane removal during cell contraction and re-insertion during apical surface expansion.

A series of ratcheted actomyosin contractions which become progressively stronger during ingression promote apical area loss of NBs [Simoes et al., 2017] (Figs. 1D,E; S1A). Notably, contraction-expansion cycles, which are ∼2.5 minutes in length, correlate with fluctuations in Crb surface levels. 79.5% of apical contractions resulted in a reduction of Crb by 13.5+/-11% each, whereas Crb levels increased by a similar degree during 76.4% of apical expansions, resulting in the maintenance of apical Crb levels during early ingression. In contrast, during late ingression, 89.6% of apical contractions reduced total apical Crb levels by 29+/-18% each, and a larger fraction of apical expansions (41.3% compared to 23.6% during early ingression) also contributed to Crb loss (Figs. 1E-G; S1A). Thus, Crb is actively removed from the membrane during contraction and secreted during expansion (Fig. 1G). The balanced decrease of Crb during contraction and increase during expansion in early ingression is consistent with normal protein turnover that maintains a uniform surface level. However, the shift to enhanced reduction of Crb during both contraction and expansion during late ingression suggests a change in the underlying mechanism of how Crb surface stability is regulated.

### Loss of Crb from the NB apical membrane promotes cell ingression

Crb stabilizes the apical membrane of epithelial cells and is lost during ingression [Tepass et al., 1990; Wodarz et al., 1995]. We overexpressed Crb to test whether its loss is required for normal ingression. Crb overexpression consistently slowed ingression rates (Figs. 2A-C), causing a 57% increase in the amplitude of apical expansions and a 32% reduction in the duration of apical contractions (Fig. 2D), response parameters that varied with the level of overexpression. To ask how persistence of Crb interferes with ingression we examined Ecad and myosin II distribution. Increasing Crb reduced Ecad levels and disrupted the apical AJ with Ecad spreading through the lateral membrane (Figs. 2E). Crb overexpression also caused a decrease in junctional myosin and an increase in medial myosin levels (Fig. 2F). Moreover, NBs in Crb overexpressing embryos displayed unstable medial myosin networks, with higher rates of medial myosin assembly and disassembly compared to controls (Fig. 2G,H). These results suggest that elevated levels of Crb delay ingression by favoring apical expansions through the apparent destabilization of junctional and medial myosin networks, which both contribute to the apical constrictions of ingressing NBs [Simoes et al., 2017].

**Figure 2.**
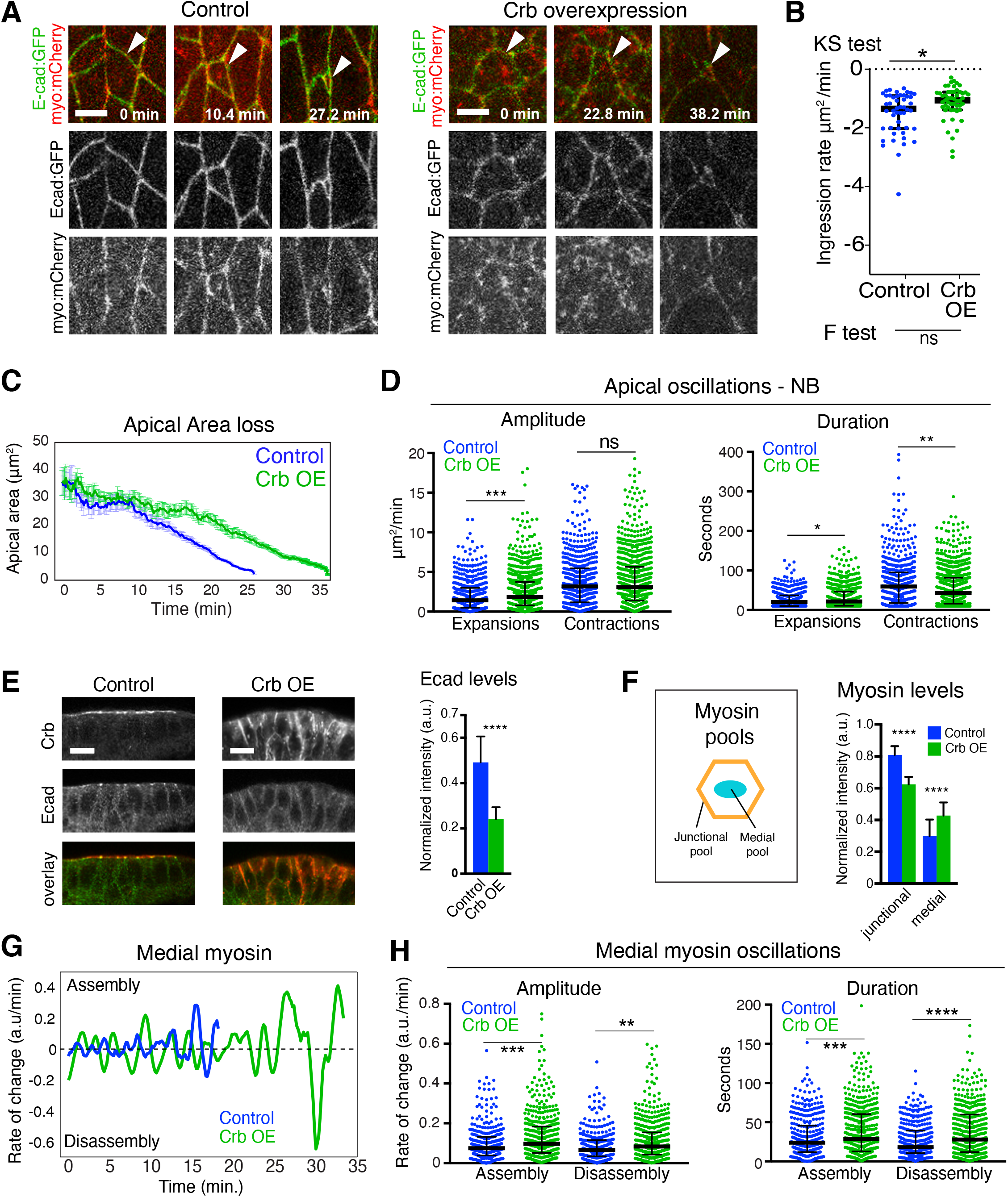
Crb overexpression delays NB ingression. **(A)** Ingressing live NBs (arrowheads) expressing endo-Ecad::GFP and sqh-Sqh::mCherry in H_2_O injected (control) and Crb overexpression (OE) embryos. T=0 min, onset of ingression. Scale bars, 5 µm. **(B)** NB ingression speed in Crb-OE (51 NBs, 10 embryos) and controls (48 NBs, 6 embryos). Median values: -1.35 (control), -1.0 (Crb-OE), KS test: * p=2.5×10^−2^; F-test (to test for differences in variance of ingression speed): ns (not significant), p=0.07; Bars are IQRs. **(C)** Apical area loss of NBs is slowed in Crb OE embryos. Control (blue line; 20 NBs, 2 embryos, 159 time points spaced by 10 sec) versus Crb OE (green line; 26 NBs, 5 embryos, 220 time points spaced by 10 sec). **(D)** Amplitude and duration of apical contractions and expansions of NB in Crb OE embryos (26 NBs, 5 embryos) versus controls (20 NBs, 2 embryos). Median amplitudes of expansions for control/Crb OE, 1.17/1.84 μm^2^/min; ****, p=5×10^−4^; and contractions, 2.76/3.04 μm^2^/min; ns, p=0.43. Median durations of expansions for control/Crb OE, 17.39/18.87 sec; *, p=0.01; and contractions, 48.64/33.06 sec, **, p =1×10^−3^ (KS test); 241–566 events per condition. **(E)** Ventral ectoderm in late stage 8 control and Crb OE embryos, stained for Crb (red) and Ecad (green). Apical is up. Note ectopic Crb on the lateral cell membrane in Crb OE cells and loss of apical Ecad signal. Scale bars, 5 µm. Quantification of Ecad levels in ingressing NBs. N values as in (D). **** p=9.6×10^−67^ (2-tailed T test; mean +/- SD). **(F)** Myosin levels in NBs of Crb OE embryos. Schematic of junctional and medial myosin pools in the apical domain of ingressing NBs. Crb OE reduces junctional myosin and and enhances medial myosin. N values as in (D). ****, p=8.2×10^−136^; 8.2×10^−38^ (2-tailed T test; means +/- SD). **(G)** Representative plots showing the rate of medial myosin change (a.u./min) during ingression in a control and a Crb OE NB. Positive rates indicate medial myosin assembly; negative rates indicate disassembly. T=0 min, onset of ingression. **(H)** Medial myosin assembly and disassembly rates (Amplitude) during ingression. Duration indicates total time medial myosin spent increasing/assembling or decreasing/disassembling. N values as in (D). Median rates of medial myosin change of control/Crb OE for assembly, 0.07/0.1 a.u./min; ***, p=3.0×10^−4^; and disassembly, 0.06/0.08 a.u./min; **, p=2.6×10^−3^. Median durations for control/Crb OE for assembly, 24/28 sec; ***, p=1×10^−3^; and disassembly, 18/28 sec, ****, p =7.9×10^−6^ (KS test); 328– 553 events per condition.

The persistence of apical Crb for much of the ingression process raises the question whether Crb plays an active role during NB ingression. Notably, NBs tended to ingress faster in Crb-depleted embryos (Figs 3A; S1B,C), although average ingression rates did not show a significant difference (Fig. 3B). However, we found a wider range in the rates of apical area loss in Crb-depleted embryos (0.2-6.2 µm^2^/min) compared to controls (0.2-3.8 µm^2^/min) (as assessed with an F-test; Fig. 3B). 25% of NBs ingressed much faster on average upon loss of Crb than controls (2.6-6.2 µm^2^/min versus 1.9-3.8 µm^2^/min, respectively) (Fig. 3B,C). NBs ingressing faster displayed a 3-fold increase in the amplitude of apical contractions, which also lasted 38% longer than in slower NBs (Fig. S1D). These findings indicate that Crb surface levels are an important determinant of ingression dynamics and ensure that NBs across the neuroepithelium complete ingression in a narrow time window.

**Figure 3.**
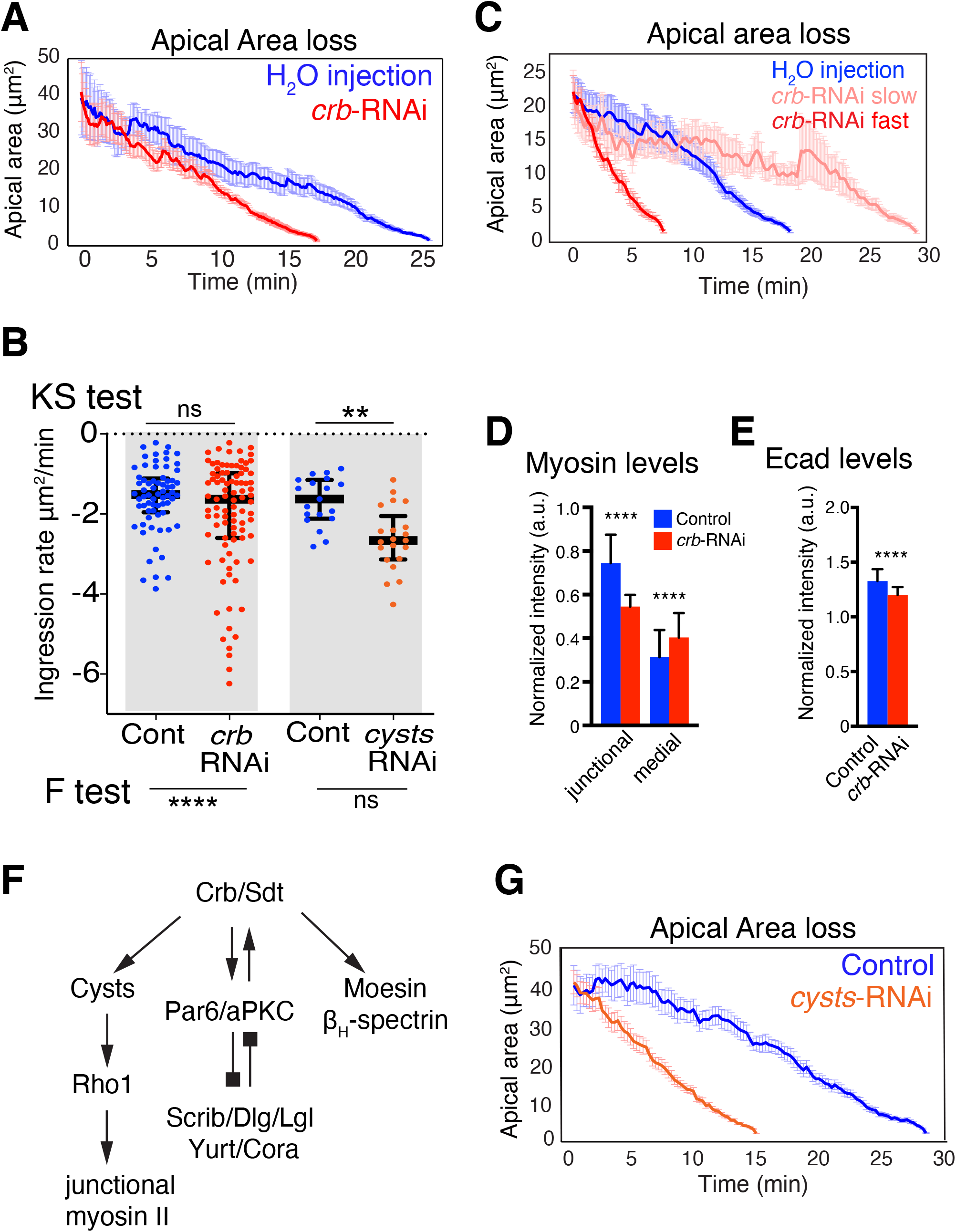
Crb and the RhoGEF Cysts are required for normal NB ingression dynamics. **(A)** Mean apical area loss during NB ingression in *crb* dsRNA injected embryos. T=0 min, onset of ingression. Means +/- SEM. Control: 20 NBs, 3 embryos, 260 time points spaced by 6 sec, *crb*-RNAi: 38 NBs, 5 embryos, 176 time points spaced by 6 sec. **(B)** NB ingression speed in *crb*-RNAi (n=91 NBs, 12 embryos), *cyst*-RNAi (21 NBs, 3 embryos), and control embryos (68 NBs, 10 embryos; 19 NBs, 4 embryos, respectively). Median values: -1.5 (H_2_O injected), -1.6 (*crb*-RNAi), -1.6 (*cysts*-RNAi control), -2.6 (*cysts*-RNAi). KS tests: ns, p=0.09, ** p=0.0014; F-tests (to test for differences in variance of ingression speed): **** p=1.7×10^−5^; ns, p=0.23. Bars are IQRs. **(C)** Mean apical area loss, the 10 fastest and the 10 slowest ingressing NBs in *crb*-RNAi (5 embryos; control, H_2_O injected, 20 NBs, 3 embryos). T=0 min, time point when the average apical surface of each ingressing cell population was 22 µm^2^. Means +/- SEM. **(D-E)** Junctional and medial myosin levels (D, ****, p=2.5×10^−85^; 3.4×10^−19^;) and Ecad (E, **** p=1.1×10^−40^) in ingressing NBs from *crb* dsRNA injected and H_2_O-injected embryos. N values as in (B). (2-tailed T test). Means +/- SD. **(F)** Cysts is one of several Crb complex (Crb/Sdt) effectors. Cysts stimulates Rho1 to enhance junctional myosin II. Other Crb/Sdt effectors are the Par6/aPKC complex which undergoes negative feedback regulation with basolateral polarity proteins (Scrib/Dlg/Lgl and Yurt/Cora), and Moesin and β_H_-Spectrin which support apical membrane stability [Tepass, 2012; Silver et al., 2019]. **(G)** Mean apical area loss during NB ingression in *cysts-*RNAi embryos compared to best-match controls. T=0 min indicates the onset of ingression. Data presented are means +/- SEM. N values: control (19 NBs, 4 embryos, 97 time points spaced by 15 sec) versus *cysts*-RNAi (21 NBs, 3 embryos, 59 time points spaced by 15 sec).

Loss of Crb resulted in a significant decrease in the amount of junctional myosin in ingressing cells, and also led to a moderate increase in medial myosin levels compared to controls (Figs. 3D; S1C). Both, fast and slow NBs showed enhanced rates of medial myosin assembly compared to controls, but in slow cells medial myosin was less stable due to higher disassembly rates (Fig. S1E,F). In addition, Crb-depleted NBs had reduced Ecad (Fig. 3E), with fast ingressing cells having significantly lower Ecad levels than slow cells (Fig. S1G). Thus, reduction in Ecad and junctional myosin levels, and greater variation in medial myosin disassembly rates may cause the enhanced variability in ingression dynamics in embryos lacking Crb.

To test whether reduction of junctional myosin associated with loss of Crb is responsible for the differences in NB ingression dynamics we examined NBs in embryos depleted of the RhoGEF Cyst. Crb recruits Cyst to the apical junctions where Cyst stimulates the Rho1-Rho kinase-myosin II pathway (Fig. 2F) [Silver et al., 2019]. Loss of Cyst reduces junctional myosin in the neuroepithelium to a similar degree as observed in Crb-depleted embryos [Silver et al., 2019; Garcia De Las Bayonas et al., 2019]. Thus, analysis of Cyst allowed us to isolate the impact Crb has on myosin from its other interactions with the Par6/aPKC complex, Moesin, or β_H_-Spectrin (Fig. 2F) [Tepass, 2012]. NB ingression speed in embryos lacking Cyst was consistently faster than controls, and did not show the variability observed upon the loss of Crb (Fig. 2B,G). Thus, isolating the impact of Crb on junctional myosin from other Crb activities through removal of Cyst revealed that the Crb-mediated stabilization of junctional myosin reduces ingression speed.

Ingression speed and the periodicity of apical contractions of NBs depend on the resistance and pulling forces exerted by neighboring non-ingressing cells (NICs) as NICs themselves undergo oscillating actomyosin contractions [Simoes et al., 2017]. Like NBs, NICs in Crb-depleted embryos display higher levels of medial myosin and lower levels of junctional myosin compared to controls [Silver et al., 2019], raising the possibility that defects in force balance between NBs and NICs could contribute to increased variation in ingression speed between NBs. We tested this hypothesis by comparing junctional and medial myosin pools between NBs and their surrounding NICs. In controls, the NB/NIC ratios of junctional or medial myosin levels positively correlated with ingression speed. In contrast, ingression speed in Crb-depleted embryos only positively correlated with medial but not junctional NB/NIC myosin ratios (Fig. S2).

Taken together, our findings support the view that a balance of NB intrinsic and extrinsic forces affect ingression speed. Because the reduction of junctional myosin seen in NBs and NICs upon loss of Crb or Cyst accelerates ingression in many NBs rather than slowing it down, we suggest that the reduced junctional myosin in NICs dominates the impact on ingression, reducing the resistance to apical constriction of NBs, thereby causing ingression in many NBs to speed up. The greater variability in ingression speed seen with the loss of Crb compared to the loss of Cyst may indicate that additional Crb functions contribute to the regulation of NB ingression. Thus, Crb persistence during early ingression with normal protein turnover rates ensures robust ingression dynamics and ingression in a normal timeframe.

### Crb ubiquitination-driven endocytosis is essential for NB ingression

One striking observation during late ingression was the enrichment of prominent Crb-positive cytoplasmic puncta within NBs (Fig. 4A,C). These puncta were mature endosomes as colocalization was observed with the endosomal markers Hrs, Vps26, Rab5, and Rab7 among others (Figs. 4B,D; S3), suggesting that Crb is removed from the plasma membrane through endocytosis. These endosomes were also enriched in Ecad and the Notch ligand Delta, which signals to surrounding NICs resulting in an epidermal fate [Hartenstein and Wodarz, 2013] (Fig. 4B,D). Crb-positive endosomes persisted in NBs for some time after ingression was completed (Figs. 4A; S3Q,R).

**Figure 4.**
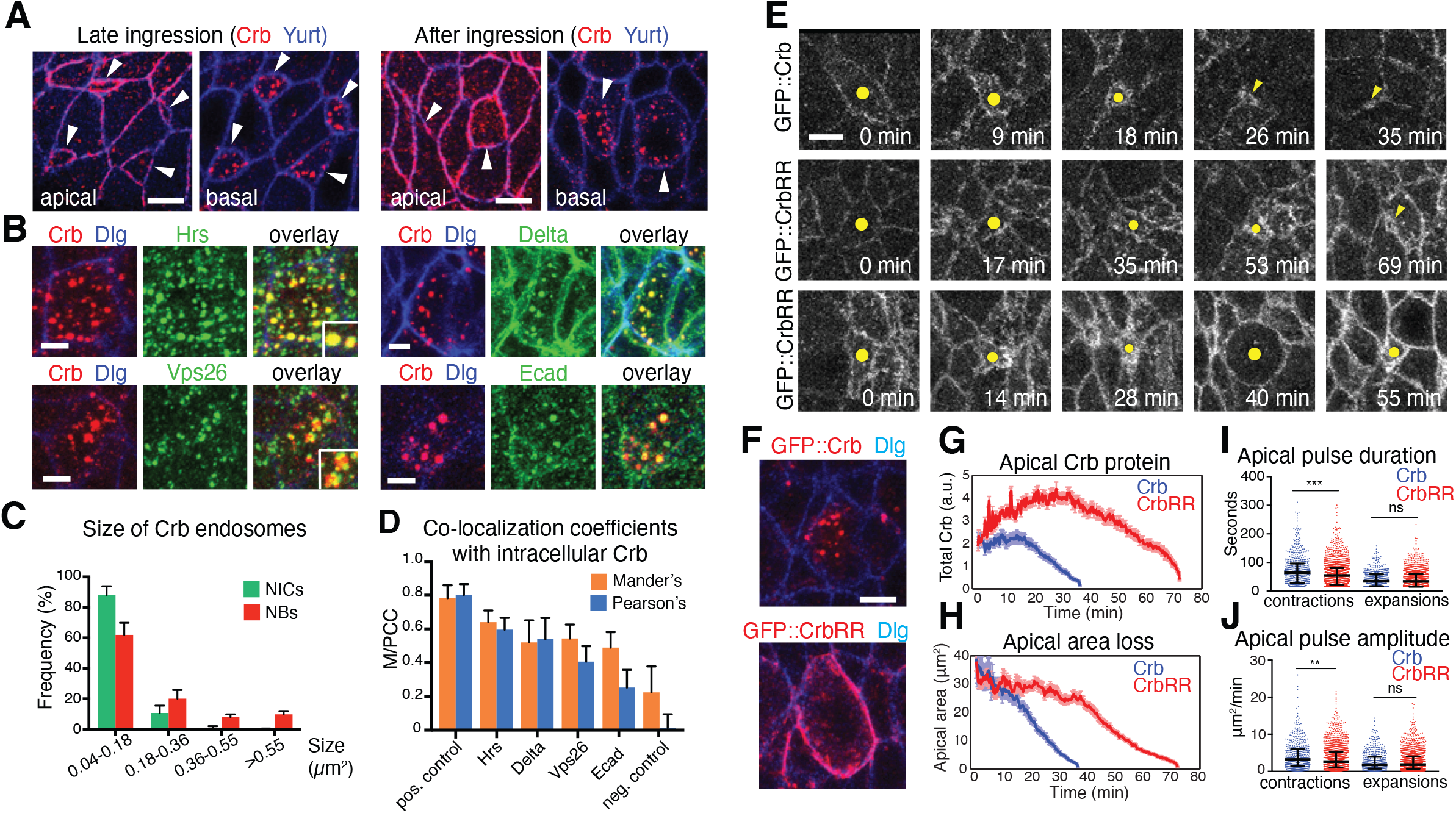
Crb endocytosis facilitates NB ingression. **(A)** In late ingressing NBs, Crb is found at the apical membrane (apical, arrowheads) and in large endosomes 0.9 µm below the apical surface (basal, arrowheads). After ingression, Crb is seen in large endosomes in the perinuclear region. Arrowheads in apical sections point to the position of NBs prior to ingression. Arrowheads in the basal sections point to NBs 6 µm below the surface. Yurt labels the basolateral membrane. Scale bars, 5 µm. **(B)** Single ingressed NBs displaying co-localization of Crb and the endosome markers Hrs, Vps26, Delta, and Ecad. The plasma membrane is labelled with Discs Large (Dlg). Scale bars, 2.5 µm. **(C)** Frequency of Crb-positive endosomes in NBs after ingression versus adjacent non-ingressing cells (NICs) according to endosomal size. Ingressed NBs contain larger Crb-positive endosomes than NICs. N=2024 endosomes from 84 NBs and 751 endosomes from 77 NICs (5 embryos). **(D)** Mander’s and Pearson’s co-localization coefficients (M/PCC) of intracellular Crb and markers shown in (B) in ingressed NBs. See Fig. S3 for positive and negative co-localization controls. 67-109 NBs from 3-7 embryos analyzed per condition. **(E)** Stills from time-lapse movies of ingressing NBs expressing GFP::Crb (control) or GFP::CrbRR (dots and arrowheads). Control NBs lost their apical domain within ∼35 min (35 NBs, 6 embryos); GFP::CrbRR expression strongly delayed ingression: 75% of NBs completed ingression after ∼70 min (middle panels; 55 NBs, 6 embryos) and 25% divided on the embryo surface (lower panels; 18 NBs, 6 embryos). Note NBs at 40 minutes showing mitotic rounding. Scale bar, 5 µm. **(F)** NBs at stage 9 expressing GFP::Crb (84 NBs, 4 embryos) or GFP::CrbRR (115 NBs, 7 embryos) in the absence of endogenous Crb (*crb*^*11a22*^ mutant embryos). GFP::CrbRR is mostly retained cortically (95/115 NBs), while GFP::Crb localizes to endosomes (84/84 NBs). Scale bar, 5 µm. **(G)** Total GFP::Crb and GFP::CrbRR fluorescence levels at the junctional domain during ingression. N values as in (F). Means +/- SEM. **(H)** Mean apical area loss during ingression in GFP::Crb (35 NB, 6 embryos) or GFP::CrbRR (55 NBs, 6 embryos) expressing NBs. T=0 min indicates the onset of ingression. Data presented are means +/- SEM. **(I**,**J)** Duration and amplitude of individual apical contractions/expansions during ingression in GFP::Crb versus GFP::CrbRR expressing NBs. Median duration for GFP::Crb/GFP::CrbRR of contractions, 64.1/54.6 sec, ***, p=1×10^−4^; expansions, 34/33.6 sec, ns (not significant), p =0.51 (KS test); Median amplitudes for GFP::Crb/GFP::CrbRR (of contractions, 3.2/2.6 μm^2^/min; **, p=1×10^−3^; expansions, 1.7/1.8 μm^2^/min; ns, p =0.94 (KS test); n =514–1553 events per condition.

How does Crb endocytosis impact NB ingression? To address this question we generated a form of Crb that showed strongly reduced endocytosis. The 37 amino acid cytoplasmic tail of Crb contains two Lysine (K) residues. K residues are potential targets for ubiquitination, which signals endocytosis [Haglund and Dikic, 2012]. K by Arginine (R) exchange to block potential ubiquitination of Crb (GFP::CrbRR) resulted in an excessive apical accumulation of GFP::CrbRR during both early and late ingression (Fig. 4E,G), and its persistence in the NB plasma membrane well after ingression (95/115 S1 NBs at stage 9), rather than a translocation into endosomes as seen with GFP::Crb (84/84 NBs; Fig. 4F). GFP::CrbRR also accumulated at the cortex of NICs, which caused an excessive apicalization as was reported for Crb overexpression [Wodarz et al., 1995]. These results suggest that ubiquitination of Crb is an important signal for its internalization.

GFP::CrbRR expression caused a failure of ingression in ∼25% of NBs, marked by a NB-type division within the neuroepithelium, which normally occurs after ingression below the epithelium (Fig. 4E-bottom panels; Sup. Videos 1 and 2). ∼ 75% of NBs ingressed successfully but took twice as long to complete ingression (Fig. 4E-middle panels, H; Sup. Videos 1 and 3). Blocking Crb endocytosis also reduced the duration and amplitude of apical contractions in ingressing NBs by 19% and 15%, respectively, relative to control NBs expressing GFP::Crb (Fig. 4I,J). These results indicate that Crb ubiquitination and subsequent endocytosis are crucial for apical membrane dynamics during early and late NB ingression, and that Crb ubiquitination is a key molecular determinant of normal ingression.

### Endocytosis, degradation, and recycling regulate NB ingression

To better understand how endocytosis contributes to the removal of the apical domain during NB ingression, we tracked membrane internalization using the lypophilic dye FM4-64 [Rigal et al., 2015] and quantified its incorporation into newly formed endosomes (Fig. 5A,B). Irrespective of injecting FM4-64 into the perivitelline space or inside the embryo (facing the apical or basal side of the neuroepithelium, respectively) newly formed endosomes originated within 1.0-6.5 μm below the apical plasma membrane (total cell depth: ∼30 μm), indicating that endocytosis predominantly occurs apically (Sup. Video 4). Interestingly, we found that the density of apical FM4-64 positive vesicles increased in NBs during late ingression but remained constant in NICs (Fig. 5A,B; Sup. Video 5). These findings are consistent with an enhanced accumulation of GFP::Dynamin and GFP::Clathrin, two endocytotic markers [Mettlen et al., 2018], at the NB apical domain compared with NICs (Fig. 5C-F). We conclude that the enhanced apical contractions observed during NB ingression [Simoes et al., 2017; An et al., 2017] correlate with an increase in apical endocytosis.

**Figure 5.**
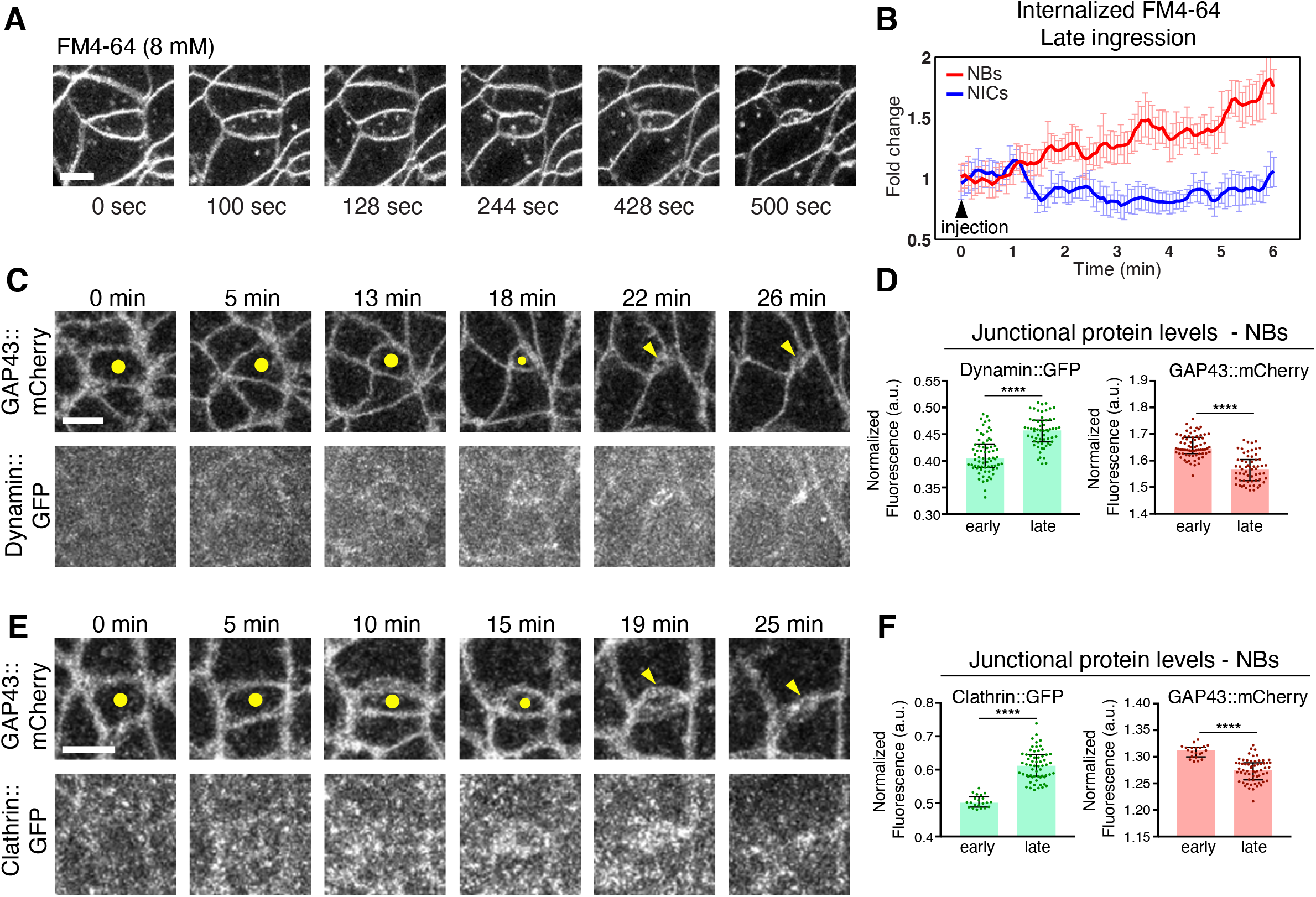
Upregulation of endocytic regulators and apical membrane internalization during NB ingression. **(A-B)** NBs increase apical endocytosis. Ingressing live NB (cell in the center; A) after injection of the lipophilic, non-cell permeable fluorescent dye FM4-64 at 8 mM, which marks the plasma membrane and apical vesicles (endosomes). Scale bar, 5 µm. (B) Intracellular FM4-64 in late ingressing NBs (14 NBs, 4 embryos) and temporally matched non-ingressing cells (NICs; 14 cells from 4 embryos), normalized to the initial time point of movie recording (∼2 minutes after injection of the dye into the perivitelline space of the embryo). The initial mean apical area of NBs at T=0 min was ∼19 µm^2^. Over the course of 6 minutes, the average fluorescence of internalized FM4-64 increased in ingressing NBs and kept constant in NICs. T=0 min, start of video recording. Means +/- SEM. **(C-F)** Ingressing live NB co-expressing Dynamin::GFP and GAP43::mCherry (C,D) or Clathrin::GFP and GAP43::mCherry (E-F) Dots/arrowheads indicate the NB apical domain. T=0 min, onset of ingression. Scale bar, 5 µm. Normalized fluorescence intensity during early and late ingression. Individual dots are averages of protein levels at the cell junctions of 13 temporally registered ingressing NBs from 2 embryos (D) or 31 temporally registered ingressing NBs from 7 embryos (F). Bars are medians and error bars are IQR.

We next reduced endocytosis by knockdown of the endocytic adaptor AP2α, which interacts with Crb [Lin et al., 2015], or by using a thermosensitive Dynamin allele (Dyn^TS^). AP2α depletion reduced the amplitude of apical contractions in NBs by 34% whereas amplitude of expansion and the duration of contractions and expansions remained normal (Fig. 6A,B). This reduced ingression speed (Fig. 6A-bottom panels), or prevented ingression of 38% of NBs, which divided on the embryo surface (Fig. 6A-middle panels; Sup. Videos 6-8). Dyn^TS^ embryos grown at the restrictive temperature from the onset of ingression showed a delay in apical domain loss (Figs. 6C; S4). 56% of NBs constricted apically but failed to complete ingression and 22% of NBs divided within the neuroepithelium. Blocking Dynamin also resulted in elevated apical Crb and Ecad concentrations in ingressing NBs (Figs. 6C; S4). Conversely, we increased apical endocytosis through the expression of a constitutively active form of the early endocytic regulator Rab5 (Rab5-CA). This accelerated the rate of apical membrane removal 1.8-fold, increased the amplitude of apical contractions by 38% and decreased the duration of apical expansions by 21%, relative to controls (Fig. 6D). Together, these results highlight a pivotal role for early endocytic regulators in directing Crb internalization and in regulating the ratcheted contractions that drive NB apical domain loss.

**Figure 6.**
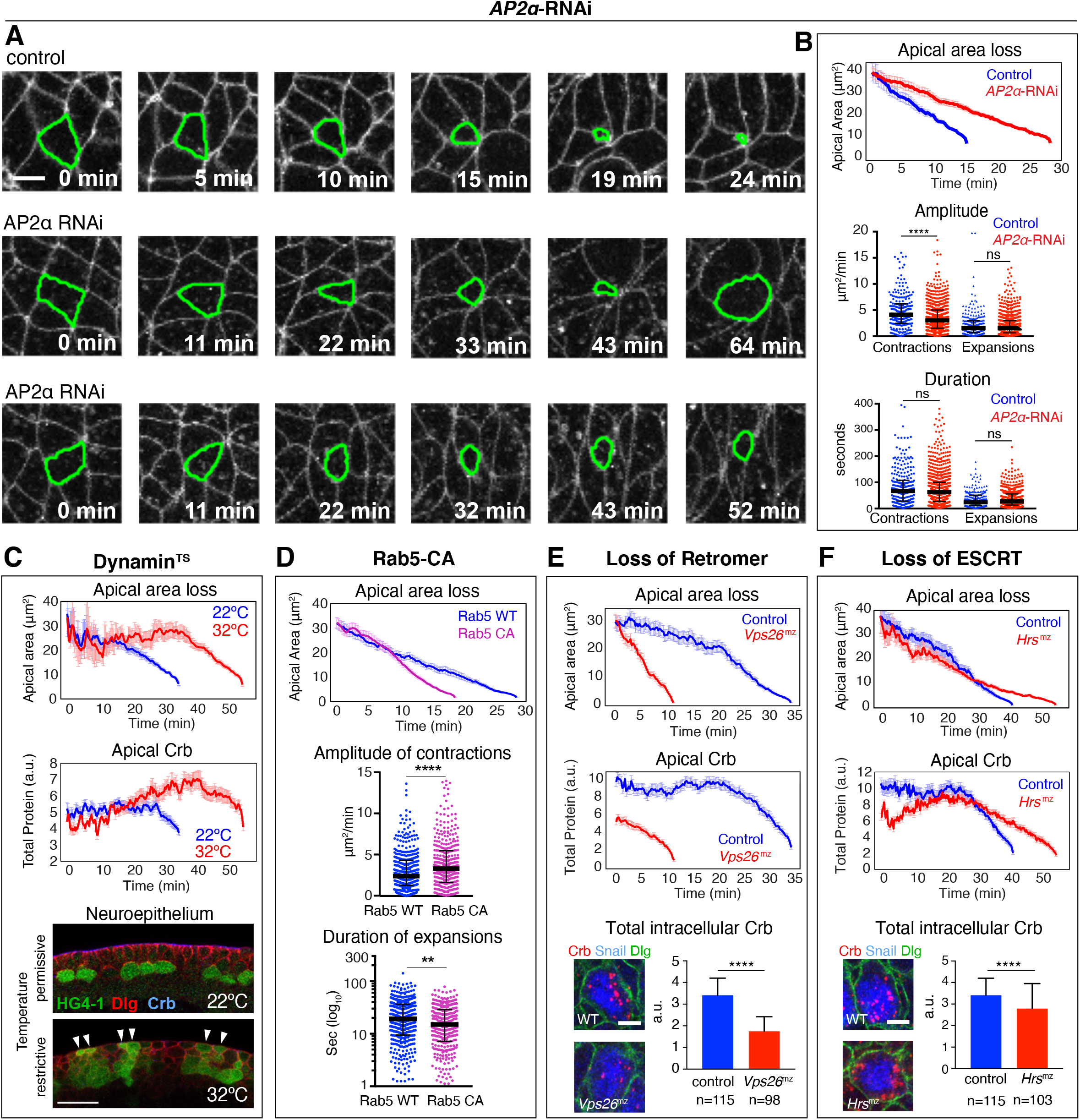
Endocytosis and Vesicle trafficking are essential for NB ingression. **(A-C)** Loss of AP2 and Dyn function blocks endocytosis. (A) Whereas control NBs lose their apical domain within ∼24 min (n=40 NBs, 2 embryos), knockdown of AP2α (*AP2α*-RNAi) results in significant delays in ingression (n=87 NBs, 4 embryos) or incomplete ingression, with 24% of NBs dividing on the embryo surface (middle panels) and 14% of NBs apically constricted but failing to complete ingression (lower panels). Membrane labeled with GAP43::mCherry, Scale bar, 5 µm. (B) Quantification of ingression dynamics of NBs in *AP2α*-RNAi (n=66 NBs, 3 embryos) versus control embryos (n=40 NBs, 2 embryos). *AP2α*-RNAi NBs show slower apical area loss and reduced amplitude of contractions compared to control. Control cells were only tracked until their apical domain reached 9 µm^2^ (in contrast to ∼1.5 µm^2^ in other figures) as 38% of NBs in *AP2α*-RNAi only constricted their apices to that value, but not below. Median amplitudes of contractions for control or *AP2α*-RNAi, 4.1/3.0 μm^2^/min, **** p=4×10^− 6^, and expansions, 1.5/1.5 μm^2^/min, ns, p=0.88 (KS test). Median durations of contractions for control or *AP2α*-RNAi, 67.2/62.7 sec; ns, p=0.22, and expansions, 24/27.4 sec, ns, p =0.27 (KS test); 233–855 events per condition. (C) Mean apical area loss and total apical Crb levels during NB ingression in Dyn^TS^ (*shibire*^*ts1*^ mutant) embryos at 22ºC and 32ºC (32 NBs from 3 embryos at 22ºC and 31 NBs from 2 embryos at 32ºC). Lower panels show cross-sections of ventral ectoderm of stage 10 embryos expressing the NB reporter HG4-1 at the permissive temperature (22ºC) or restrictive temperature (32ºC) where some NBs failed to ingress (arrowheads). N=8 and 11 Dyn^TS^ embryos at 22ºC and 32ºC, respectively. Apical side, up. Scale bar, 15 µm. T=0 min, onset of ingression; means +/- SEM. Rab5 activity promotes ingression. Mean apical area loss during NB ingression in embryos expressing constitutive active Rab5-CA (58 NBs, 4 embryos) or normal Rab5 (Rab5-WT; control; 36 NBs, 3 embryos). T=0 min, onset of ingression; means +/- SEM. Amplitude and duration of individual apical contractions or expansions during ingression in Rab5-WT versus Rab5-CA embryos. Median amplitude of contractions for Rab5-WT or Rab5-CA: 2.4/3.3 μm^2^/min; ****, p=4.8×10^−5^. Median duration of expansions for Rab5-WT or Rab5-CA: 19/15 sec (log_10_), **, p= 1.4×10^−3^(KS test); 341-448 events per condition. **(E)** Retromer limits ingression speed. Mean apical area loss during NB ingression and total apical Crb levels in *Vps26*^*mz*^ mutants (23 NBs, 5 embryos; mz indicates maternal-zygotic mutants) versus controls (59 NBs, 12 embryos). Total intracellular levels of Crb in ingressed NBs from *Vps26*^*mz*^ mutants (98 NBs, 7 embryos) and controls (115 NBs, 5 embryos). **(F)** ESCRT enhances loss of apical Crb and apical membrane. Mean apical area loss during NB ingression and total apical Crb levels in *Hrs*^*mz*^ mutants (36 NBs, 7 embryos) versus controls (40 NBs, 5 embryos). Total intracellular levels of Crb in ingressed NBs from *Hrs*^*mz*^ mutants (103 NBs, 6 embryos) and controls (115 NBs, 5 embryos). In (E) and (F): Snail (blue) identifies NBs. Scale bar, 5 µm. T=0 min, onset of ingression; means +/- SEM.

Endosomal Crb colocalizes with Hrs, a component of the ESCRT complex which facilitates late endosomal/lysosomal processing and protein degradation [Vietri et al., 2020], and Vps26, a component of the Retromer complex that is crucial for maintaining Crb surface levels in the neuroepithelium through recycling [Zhou et al 2011; Pocha et al., 2011; McNally and Cullen, 2018]. We therefore assessed whether ESCRT or Retromer function is required for NB ingression. Strikingly, loss of Retromer function, which reduced Crb levels by half both apically and intracellularly in NBs, strongly accelerated ingression by 67% (2.5+/-0.9 µm^2^/min versus 1.5+/- 0.9 µm^2^/min in controls) (Fig. 6E). This shows that recycling is crucial for maintaining Crb surface levels in NBs similar to the neuroepithelium as a whole, and indicates that Retromer-mediated recycling counteracts the loss of the apical domain. The faster ingression seen with the loss of Retromer function compared to the loss of Crb (Fig. 6E versus Fig. 3A) suggests that the Retromer complex recycles additional factors that stabilize the apical domain of NBs. Ecad, which colocalizes with Crb in NB endosomes (Fig. 4B,D) is an attractive candidate. In contrast to Retromer, blocking the ESCRT complex by removing Hrs function had no major impact on ingression dynamics. NBs tended to ingress slower in these embryos, a delay that may be specific to late ingression (Fig. 6F), when Crb and other transmembrane proteins are normally rapidly translocated from the apical domain into an ESCRT-positive endocytic compartment. NBs in embryos that lack ESCRT function retained apical Crb 15 minutes longer than control NBs and displayed reduced intracellular Crb accumulation in a more diffuse pattern than controls (Fig. 6F). We conclude that ESCRT and Retromer machineries are important determinants of NB ingression dynamics.

### Neuralized resolves the Crb-Sdt complex for rapid internalization of Crb during late ingression

Our analysis suggests that the rapid loss of Crb from the NB apical membrane during late ingression is crucial for delamination. This raises the question of how Crb can deviate from steady-state turn-over to be completely removed from the membrane. In embryos lacking the Crb-binding partner Sdt, Crb is unstable and rapidly endocytosed [Tepass and Knust, 1993]. We confirmed these results with live-imaging of Crb::GFP in embryos lacking Sdt. Loss of Sdt rendered Crb mostly endocytotic across the neuroepithelium. In contrast, non-endocytosable GFP::CrbRR remained at the plasma membrane in Sdt-depleted embryos (Fig. S5A). Loss of Sdt increased late ingression speed by two-fold, while having no net effect on early ingression (Fig. 7A). We conclude that destabilization of the Crb-Sdt interaction promotes Crb endocytosis and apical domain loss specifically during late ingression.

**Figure 7.**
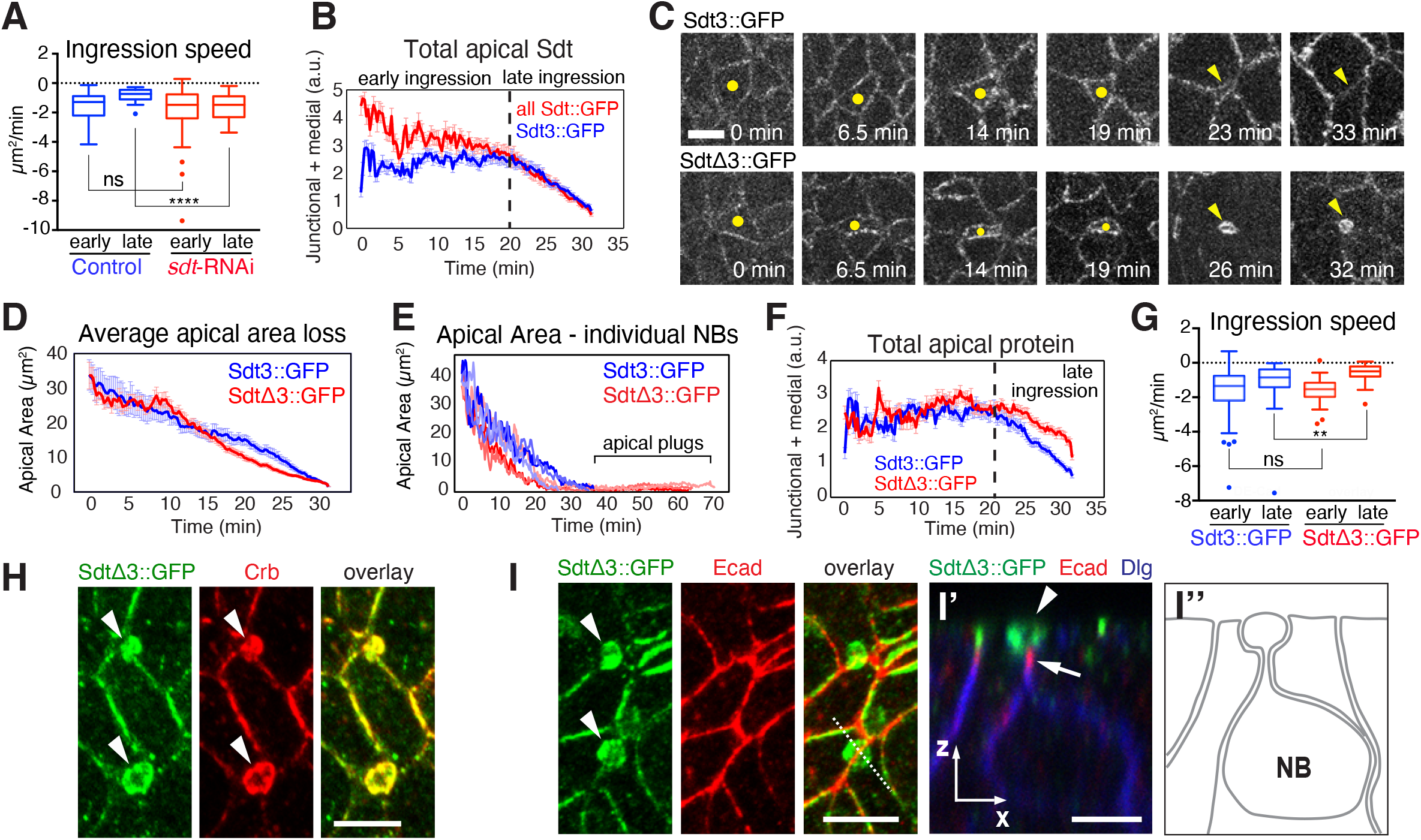
Neur-dependent disruption of the Crb complex is essential for NB ingression. **(A)** Loss of Sdt slows late ingression. Tukey’s box-and-whisker plots of NBs ingression speed (apical area reduction) in *sdt*-RNAi and controls embryos during early and late ingression. Medians (µm^2^/min) for control and *sdt*-RNAi: -1.3/-1.6 µm^2^/min (early ingression), ns, p= 0.44 and -0.6/-1.3 µm^2^/min (late ingression), ****, p=1.9×10^−5^ (KS test). *sdt-*RNAi: 54 NBs, 6 embryos; controls: 30 NBs, 4 embryos. **(B)** Quantification of total apical fluorescence levels for all Sdt isoforms (Sdt::GFP, 36 NBs, 6 embryos) and for Sdt3 (Std3::GFP, 48 NBs, 6 embryos). T=0 min, onset of ingression; means +/- SEM. **(C)** Ingressing live NBs in Sdt3::GFP (control) and SdtΔ3::GFP embryos (dots, arrowheads). Sdt3::GFP NBs lost their apical domain within ∼33 min (top panels, 48 NBs, 4 embryos). Apical plugs formed in SdtΔ3::GFP ingressing NBs (arrowheads in lower panels; 48 NBs, 4 embryos). Scale bar, 5 µm. **(D)** Mean apical area loss during NB ingression in Sdt3::GFP and SdtΔ3::GFP embryos. N values as in (C). T=0 min, onset of ingression; means +/- SEM. **(E)** Quantification of apical area loss in 3 individual NBs from Sdt3::GFP (blue lines) and SdtΔ3::GFP (red lines) embryos during ingression. SdtΔ3::GFP NBs maintain an apical plug for 30-40 minutes after control cells have completed ingression. T=0 min, onset of ingression. **(F)** Quantification of total Sdt3::GFP and SdtΔ3::GFP fluorescence at the apical domain during ingression. n values as in (C). T=0 min, onset of ingression.; means +/- SEM. **(G)** SdtΔ3::GFP embryos show reduced late ingression speed. Tukey’s box-and-whisker plots of ingression speed in Sdt3::GFP and SdtΔ3::GFP NBs during early and late ingression. Medians for Sdt3::GFP and SdtΔ3::GFP: -1.4/-1.3 µm^2^/min (early ingression), ns, p= 0.12 and -0.8469/-0.4946 µm^2^/min (late ingression), **, p= 1×10^−3^(KS test); n values as in (C). **(H)** Co-localization of SdtΔ3::GFP and Crb at apical plugs in stage 9 SdtΔ3::GFP embryos (arrowheads). Scale bar, 5 µm. **(I)** Apical plugs in SdtΔ3::GFP embryos (arrowheads) do not contain Ecad. Dashed line indicates position of ZX cross-section through an apical plug shown in the right panel (I’) and schematic (I’’). The apical plug containing SdtΔ3::GFP (arrowhead) is positioned apically to Ecad/AJs (arrow). Dlg labels basolateral membrane. Scale bars, 5 µm.

Endocytosis of Crb is fostered by the E3 Ubiquitin ligase Neuralized (Neur) via interaction with an isoform of Sdt containing sequences encoded by its exon 3 (Sdt3) [Perez-Mockus et al., 2017a]. Interestingly, overactivation of Neur disrupts epithelial polarity through a deactivation of Sdt, precipitating the loss of Crb [Chanet and Schweisguth, 2012; Perez-Mockus et al., 2017a]. Sdt3 is one of multiple Sdt alternative splice forms [Bulgakova et al., 2010; Perez-Mockus et al., 2017a]. Quantitative imaging of total endogenous Sdt (Sdt::GFP) and Sdt3::GFP, a construct where only the endogenous Sdt3 isoform is GFP-tagged, revealed that Sdt3 represented ∼79% of the total apical Sdt during early ingression but was the only Sdt isoform present in NBs during late ingression (Fig. 7B). Thus, we explored the possibility that the Sdt3-Neur interaction could mediate Crb endocytosis and apical domain loss during late NB ingression.

NBs from embryos expressing Sdt3::GFP completely lost their apical domain within ∼30 minutes as in normal embryos (Fig. 7C; Sup. Video 9). Also NBs in embryos expressing an isoform of Sdt lacking exon 3 (SdtΔ3::GFP), which abrogates the interaction between Sdt3 and Neur [Perez-Mockus et al., 2017a], constricted their apical domain in a normal timeframe (Fig. 7D). However, SdtΔ3::GFP NBs failed to fully internalize the apical membrane and retained small ‘apical plugs’ that persisted for an extended period of time. These apical plugs were spherical, with a total surface of 6.2+/-3 µm^2^ representing ∼15% of the initial apical cell surface and remained detectable for at least 40 minutes beyond normal ingression time (Fig. 7C,E; Sup. Video 10). SdtΔ3::GFP showed similar apical protein levels to Sdt3::GFP at early ingression but was less efficiently removed from the apical cortex during late ingression (Fig. 7F), and apical plugs in SdtΔ3::GFP embryos accumulated high levels of SdtΔ3::GFP and Crb (Fig. 7H). Ecad was confined to a narrow neck below the plug (Fig. 7I) as the main cell body of the NB moved below the epithelium, and AJs disassembled shortly thereafter. Apical plugs remained at the embryo surface suggesting that they detached from the main cell body of the NBs. Plugs where ultimately resolved. The abnormal accumulation of apical Crb and SdtΔ3::GFP correlated with a deceleration in apical area loss during late ingression relative to Sdt3::GFP controls (Fig. 7G). These findings indicate a specific defect in internalization of Crb and apical membrane loss when the Sdt3-Neur interaction is blocked, and show that the Sdt3-Neur interaction is crucial for the removal of the NB apical domain during late ingression.

Neur is part of the Notch signalling pathway,promoting endocytosis of the Notch ligand Delta and, consequently, Notch activation and suppression of NB fate in NICs that surround a NB [Perez-Mockus and Schweisguth, 2017; Kovall et al., 2017]. Loss of Delta, Notch or Neur causes the formation of supernumerary NBs that ingress as clusters rather than as individual cells [Hartenstein and Wodarz, 2013; Simoes et al., 2017]. To validate that internalization of Crb was in part mediated by Neur, we examined Neur-depleted embryos in comparison to Delta-depleted embryos. Upon reaching an average apical surface of ∼15 µm^2^ (late ingression), Neur-depleted NBs displayed more apical Crb, and ingression speed significantly decreased compared to NBs in Delta-depleted embryos indicating that Neur acts independent of Delta in disrupting the Crb complex (Figs. 8A-C, S5B). Loss of Neur led to a strong reduction in oscillations of apical contractions in NBs likely causing the delay in ingression (Fig. 8D). To directly test whether Neur promotes Crb endocytosis in ingressing NBs, we quantified total levels of endocytotic Crb in ingressed NBs from Neur-depleted embryos and SdtΔ3::GFP embryos. Both mutant backgrounds showed a significant reduction (14-26%) in total levels of internalized Crb compared to controls (Fig. 8E,F). Taken together, our findings suggest a novel function for the Sdt3-Neur interaction in driving the efficient removal of apical membrane during late NB ingression by promoting the disassembly of the Crb complex and, consequently, Crb endocytosis.

**Figure 8:**
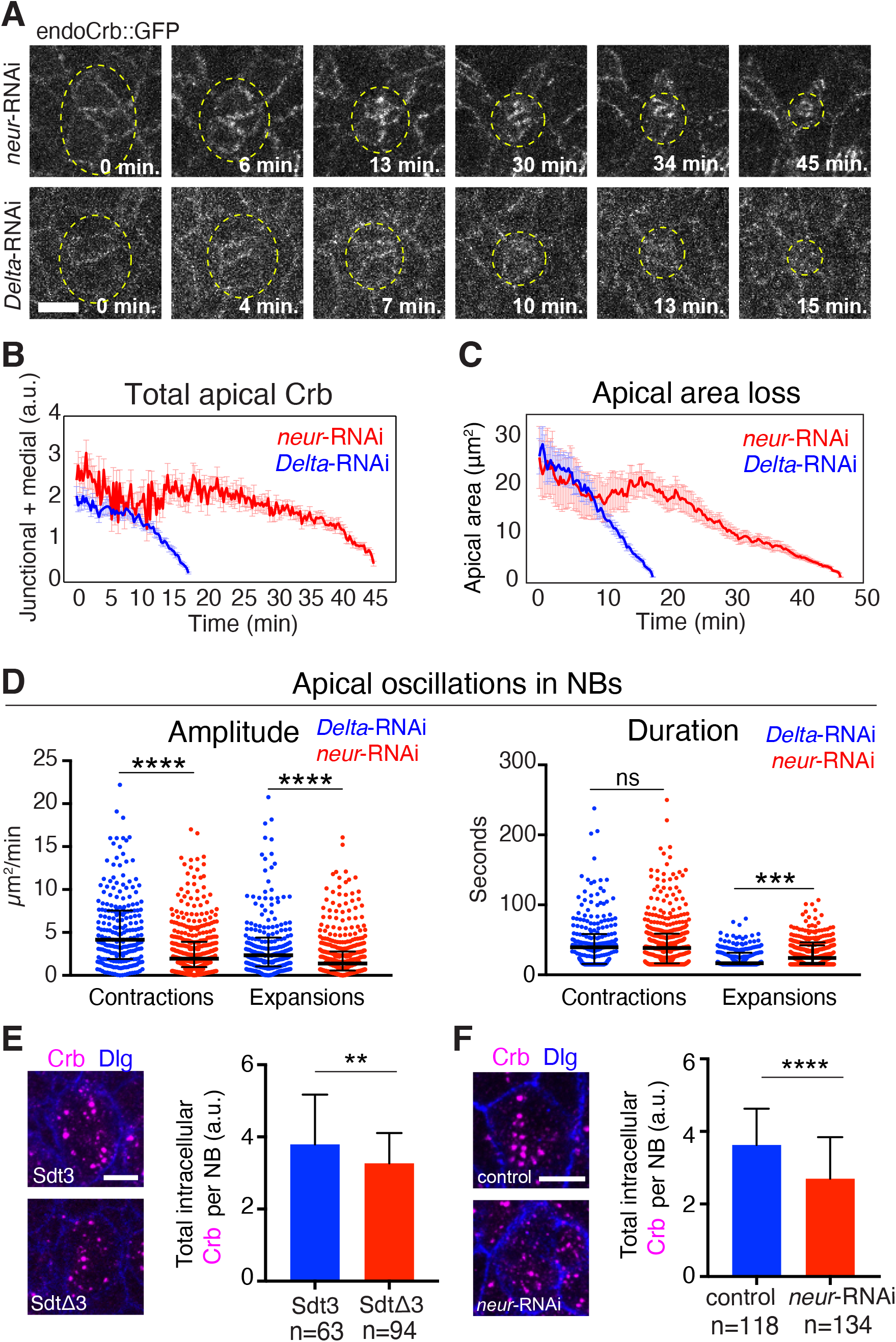
Neur, but not Delta, is required for removal of Crb from the NB apical membrane during ingression. **(A)** Endo-Crb::GFP embryos showing NBs clusters (circled) in *neur*-RNAi (20 NBs, 2 embryos) and *Delta*-RNAi (20 NBs, 2 embryos) embryos. Note ectopic accumulation of apical Crb and slower ingression with *neur*-RNAi compared to *Delta*-RNAi. Scale bar, 5 µm. **(B-C)** Total Crb::GFP (B) at apical domain (junctional and medial pools) and apical area loss (C) during ingression in *neur*-RNAi (20 NBs, 2 embryos) and *Delta*-RNAi (20 NBs, 2 embryos) embryos. *Delta*-RNAi was chosen as a control as it causes a neurogenic defect similar to *neur*-RNAi. Due to reduced levels of apical Crb in both conditions, the apical surface was traced starting at 26 µm^2^ (mid-ingression, T=0 min) in both data sets; means +/- SEM. **(D)** Amplitude and duration of apical contractions and expansions during NB ingression in *neur*-RNAi versus *Delta*-RNAi embryos. Median amplitudes for *Delta*-RNAi and *neur*-RNAi for contractions, 4.1/1.9 μm^2^/min, **** p=9.6×10^−13^; and expansions, 2.3/1.3 μm^2^/min, **** p=5.3×10^−6^, (KS test). Median durations for *Delta*-RNAi and *neur-*RNAi for contractions, 39.3/38.4 sec; ns, p=0.88; expansions, 16.6/24.1 sec, ****, p =8.2×10^−5^ (KS test); n values as in (B); 230–440 events per condition. The median amplitudes of apical contractions and expansions were reduced by 54% and 41%, respectively, in NBs of *neur*-RNAi embryos, and the median duration of apical expansions was increased by 45% compared to *Delta*-depleted controls. **(E-F)** Crb endocytosis is reduced in SdtΔ3 and *neur*-depeted embryos. Average total fluorescence of intracellular Crb per NB after ingression (stage 9) in Sdt3::GFP (63 NBs, 3 embryos) and SdtΔ3::GFP (94 NBs, 4 embryos) embryos (E). Average total fluorescence of intracellular Crb per NB after ingression (stage 9) in *neur*-RNAi (134 NBs, 7 embryos) and control (118 NBs, 6 embryos) embryos (F). Scale bar, 5 µm. Means +/- SD. **, p=3×10^−3^ ; ****, p=4.5×10^−11^ (2-tailed T test).

## Discussion

Drosophila NBs are an outstanding model for scrutinizing the cellular machineries underpinning an EMT-like process with high temporal and spatial resolution. While ingressing, a single NB sequentially loses AJs responding to tensile forces exerted by two pools of actomyosin: a planar polarized pool enriched at anterior-posterior junctions, which disassembles first, and a pulsatile pool at the free apical cortex which further tugs on shrinking junctions in a ratchet-like manner [Simoes et al., 2017; An et al., 2017]. However, while actomyosin forces reduce the apical perimeter, it remained unclear how cells lost their apical domain and how polarity regulators contribute to the dynamics of apical domain loss. We demonstrate that regulation of the Crb complex plays a key role in orchestrating apical domain loss during ingression. During early ingression, cells shrink their apical domain while retaining total levels of Crb, which is crucial for maintaining normal actomyosin in NBs and NICs to generate the tension balance required for normal ingression dynamics. During late ingression, Crb is rapidly lost from the apical domain, a process initiated by an interaction between Neur and Sdt which causes the disassembly of the Crb complex. The loss of the Crb complex then precipitates the concurrent loss of the apical membrane and AJs. A similar regulatory interplay between Crb, Sdt, and Neur was also observed during early neurogenesis in the Drosophila optic lobes [Shard et al., 2020]. Here, neural stem cells emerge from a wave front in the optic lobes rather than ingress as individual cells. Despite these topological differences, the Crb/Sdt/Neur module appears to be a common cell biological regulator of EMT during early neurogenesis.

Endocytosis and endocytic degradation and recycling are requirements for normal NB ingression dynamics. The apical membrane of neuroepithelial cells is much more active endocytically than the basolateral membrane. Notably, Crb is endocytosed during apical contractions and re-secreted during expansions, suggesting that myosin-driven cell contact contraction promotes endocytosis, consistent with recent data from mammalian cells [Cavanaugh et al., 2020], whereas expansions allow for enhanced secretion. Blocking endocytosis increases surface levels of Crb and Ecad as expected, and prevents NB ingression, whereas enhancing endocytosis accelerates ingression. Moreover, endocytic trafficking plays a key role in determining ingression speed. Loss of ESCRT complex-mediated degradation appears to enhance apical Crb and slow ingression, whereas loss of Retromer-mediated recycling dramatically reduced surface Crb and accelerated ingression. In fact, Retromer-compromised embryos showed the fastest ingression speed of any condition we have examined, suggesting that the Retromer not only recycles Crb but also other factors that counteract apical domain loss in NBs. Crb turnover during early ingression maintains a steady Crb surface abundance. During late ingression, Crb endocytosis is enhanced during both apical contraction and expansion as a result of the disruption of the Crb-Sdt interaction by Neur, which likely makes the Crb cytoplasmic tail accessible to the Clathrin adapter AP2. AP2 binds to Crb competitively with Sdt [Lin et al., 2015], facilitating the rapid endocytic removal of Crb and the apical membrane.

EMT is thought to be initiated by the expression of EMT transcription factors (EMT-TFs) of the Snail, Zeb, or bHLH families that downregulate key adhesion or polarity proteins such as Ecad and Crb [Lamouille et al., 2014; Nieto et al., 2016; Dongre and Weinberg, 2019; Yang et al., 2020]. NBs are specified through the combined action of proneural genes that include bHLH proteins of the Achaete-Scute complex (AS-C), the Snail family protein Wornui, and the SoxB family protein SoxNeuro [Hartenstein and Wodarz, 2013; Arefin et al., 2019]. However, although genes that encode Ecad and Crb are transcriptionally downregulated in NBs [Tepass et al., 1990; Tepass et al., 1996] this repression does not appear relevant for NB ingression. Replacing endogenous Ecad with a transgene expressing Ecad under the control of a ubiquitous promoter had no impact on NB ingression dynamics [Simoes et al., 2017]. Here, we show that surface levels of Crb remained high in NBs during early ingression before Crb is rapidly removed by endocytosis during late ingression. This raises the question of how the upregulation of proneural genes in presumptive NBs elicits enhanced actomyosin contractility and endocytic removal of apical membrane and junctions.

One proneural gene target is *neur* [Miller and Posakony, 2018; Arefin et al., 2019]. Neur is found throughout the neuroepithelium participating in Delta-Notch-mediated lateral inhibition to select the NB from an equivalence group of 5-7 cells [Boulianne et al., 1991; Hartenstein and Wodarz, 2013]. *neur* upregulation in ingressing NBs is thought to be part of a positive feedback that stabilizes NB fate through persistent asymmetric Delta-Notch signalling [Miller and Posakony, 2018; Arefin et al., 2019]. The increase in Neur may also be important for the effective disruption of the Crb complex to destabilize the apical domain. Neur can disrupt the Crb complex across the epithelium, but is normally prevented from doing so by Bearded proteins that act as inhibitors of Neur [Chanet and Schweisguth, 2012; Perez-Mockus et al., 2017a]. Increasing Neur concentration may overcome this inhibition in NBs. This raises the possibility that the proneural gene-dependent upregulation of *neur* contributes to the timing of ingression, consistent with our observation that in Neur-depleted embryos ingression is prolonged. Furthermore, Neur may enhance actomyosin contractility in NBs seen in late ingression [Simoes et al., 2017; An et al., 2017] as was reported for Neur in the Drosophila mesoderm [Perez-Mockus et al., 2017b]. We hypothesize therefore that Neur could be a central regulator of NB selection and ingression; stabilizing NB fate, driving apical membrane constriction through actomyosin contraction, and disrupting the Crb complex to remove apical membrane and junctions. Interestingly, we also noted that during ingression the number of alternative isoforms of Sdt is limited to Sdt3, the isoform susceptible to Neur. Hence, NBs appear to develop the molecular competence for apical membrane removal at least in part through rebalancing Sdt splice forms.

The loss of apical-basal polarity is an early event during EMT marked by the loss of epithelial AJs that can trigger expression of EMT-TFs and the disassembly of cell junctions [Ozdamar et al., 2005; Jung et al., 2019]. However, our findings indicate that the loss of apical-basal polarity in NBs is preceded by a period (∼20 min; early ingression) of ratcheted apical contractions that reduce the apical area of delaminating cells [Simoes et al., 2017; An et al., 2017]. The maintenance of normal Crb levels during early ingression is crucial for normal ingression dynamics. Crb stabilizes junctional myosin through it effector, Cyst that is recruited to the junctional domain by the Crb complex [Silver et al., 2019]. Whereas the loss of Crb or loss of Cyst causes similar reductions of junctional myosin in the neuroepithelium [Silver et al., 2019], NB ingression was consistently faster in Cyst-compromised embryos than controls. In contrast, NBs in Crb compromised embryos showed much larger variability of ingression speeds, with a small fraction of NBs ingressing rapidly while the majority was slower than controls. Thus, it is likely that Crb makes other contributions to regulating NB ingression in addition to its Cysts-mediated function in supporting junctional actomyosin. Interestingly, the mouse Crb homolog Crb2 is required for myosin organization and ingression during gastrulation [Ramkumar et al., 2016]. The predominant defect in Crb2 compromised mice appears to be a failure of ingression which may be similar to the fraction of NBs showing slower than normal ingression seen with the loss of Drosophila Crb. To what extent the differences in cell behaviour caused by the loss of Crb and Crb2 depend on the biomechanical specifics of the tissue context or result from differences in molecular pathways in which Crb and Crb2 operate remains to be explored.

## Materials and Methods

### Markers and mutants

The following fly markers and mutants were used: *w*^*1118*^ as ‘wild type’ (Bloomington Drosophila Stock Center: BDSC_3605), *endo-crumbs::GFP-C* [Huang et al., 2009], *endo-DEcad::GFP* (BDSC_60584; gift from Y. Hong, University of Pittsburgh), *y w;ubi-DEcad::GFP* [Oda et al., 2001], *sqh-sqh::mCherry* [Martin et al., 2009] and *sqh-GAP43::mCherry* (gifts from A.C. Martin, Massachusetts Institute of Technology) [Martin et al., 2010], *w FRT18E Par6*^*226*^ *P[promPar6_Par6::GFP]61-1F* [Petronczki and Knoblich, 2001], *endo-Delta::GFP* [Corson et al., 2017], *sdt::GFP, sdt::GFP3*, and *sdtΔ3::GFP* (gifts from F. Schweisguth, Institut Pasteur, Paris) [Perez-Mockus et al., 2017a], *HG4-1* (gift from Chris Q. Doe, University of Oregon) [Hirono et al., 2012], *y w;; Mi(PT-GFSTF*.*1)kst*^*MI03134-GFSTF*.*1*^ (BDSC_60193), *y w shi*^*ts1*^ (= *Dyn*^*TS*^; BDSC_7068), *y w Vps26*^*3c*^ *FRT101/FM7c* (a *Vps26* null allele; K.A.K and U.T., unpublished), *Hrs*^*D28*^*FRT40A/In(2LR)Gla, wg*^*Gla-1*^ *PPO1*^*Bc*^ (BDSC_54574), *w;; crb*^*11a22*^ *FRT82B*/*TM3 Sb* [Pellikka et al., 2002], *matαtub67-Gal4; matαtub15-Gal4* (gift of D. St Johnston, University of Cambridge, England, UK), *UAS-AP2α-RNAi* (BDSC_32866, Trip line #HMS00653, Drosophila RNAi Screening Center, Harvard Medical School), *UASp-YFP::Rab5* (BDSC_9775), *UASp-YFP::Rab5Q88L* (BDSC_9773), *w; UAS-crb*^*WT2e*^ [Wodarz et al., 1995], *w;; UASt-GFP::crb (attp2)* and *w;; UASt-GFP::crbRR (attp2), UAS-Dcr-2, w*^*1118*^ (BDSC_24646), *w; UAS-GFP-myc-2XFYVE* (BL_42712), *w; UAS-Rab7::GFP* (BDSC_42705), *w; UASt-GFP::Lamp* (BDSC_42714, gift from J. Brill, University of Toronto, Canada), *w; UAS-Rab11::GFP* (BDSC_8506), *UAS-PLCδ-PH::eGFP* (BDSC_39693), *UAS-IVS-Syn21-shi*^*ts1*^*::GFP-p10* [Pfeiffer et al., 2012], *UAS-eGFP::Clc* (BSDC_7107) (gift from T. Lecuit, Institut de Biologie du Développement de Marseille).

To overexpress Crb maternally, we analyzed the progeny of *matαtub67-Gal4 endo- DEcad::GFP/UAS-crbWT2e; sqh-sqh::mCherry/+* females crossed to *UAS-crb*^*WT2e*^ homozygous males. Control embryos were the progeny of *matαtub67-Gal4 endo-DEcad::GFP/+; sqh-Sqh::mCherry/+* females crossed to *w*^*1118*^ males. Similarly, overexpression of GFP::CrbRR and GFP::Crb was obtained by crossing *matαtub67-Gal4/+; matαtub15-Gal4/UAS-GFP::crbRR* (or *GFP::crb*) females to *UAS-GFP::crbRR* (or *GFP::crb*) males. A similar genetic scheme was employed to maternally express *UAS-GFP-myc-2XFYVE, UAS-Rab7::GFP, UAS-GFP::Lamp1, UAS-Rab11::GFP* and *UAS-PLCδ-PH::eGFP. Matαtub15-Gal4* alone was used to drive the expression of *UASp-YFP::Rab5* (Fig. S3). *Matαtub67-Gal4* recombined with *sqh-GAP43::mCherry* was used to drive the expression of *UAS-eGFP::Clc* (Fig. 5). *Matαtub15-Gal4* recombined with *sqh-GAP43::mCherry* was used to drive the expression of *UASp-YFP::Rab5, UASp-YFP::Rab5Q88L* and *UAS-IVS-Syn21-shi*^*ts1*^*::GFP-p10* (Fig. 6 and Fig. S4).

To knockdown *cysts* in the maternal germline, we crossed *Matαtub67-Gal4, endo-DEcad::GFP* (BDSC_60584) with *Df(2L)BSC301* (a deletion uncovering *cysts*; BDSC_23684); *UAS-cysts-shRNA* (BDSC_41578). Female progeny of this cross were mated with *UAS-cysts-shRNA* (BDSC_38292) males and ingression dynamics of NBs were analyzed in the resulting progeny.

To down-regulate Dynamin function and visualize NBs in fixed embryos, 0-3 hours after egg laying (AEL) Dyn^TS^ (= *shi*^*ts1*^); *HG4-1* embryos (laid at 18°C) were aged for 1.5 hours at 22°C and either kept at 22°C (permissive temperature) or transferred to 32°C (restrictive temperature) for 3 hours before fixation.

To knockdown *AP2α* during ingression, we analyzed the F2 progeny of *matαtub15-Gal4 sqh-GAP43::mCherry* females crossed to *UAS-AP2α-RNAi* males. Control embryos were generated similarly by using *w*^*1118*^ males. Females carrying *Vps26*^*3c*^ and *Hrs*^*D28*^ germline clones expressing *endo-crb::GFP* were generated using the FLP-DFS system [Chou and Perrimon, 1996]. Their respective wild-type controls expressing *endo-crb::GFP* were imaged in parallel using the same conditions.

### Immunohistochemistry

Embryos were fixed in a 1:1 mixture of 3.7% formaldehyde in phosphate buffer, pH 7.4, and heptane for 20 minutes under agitation, and devitellinized with a methanol/heptane mixture (Method A) or a 1:1 mixture of 37% formaldehyde and n-heptane for 5 minutes, followed by hand-devitellization (Method B). Method A was used in Fig. 4A, Fig. 4B (to detect Delta::GFP); Fig. 4F; Fig. 6C, E, and F; Fig. 7H, I; Fig 8E; Fig. 2E; Fig. S3A-D, F-J, L, and M. Method B was used in Fig. 4B (to detect Hrs, Vps26 and Ecad::GFP); Fig. 8F; Fig S1B; Fig. S3E, K, and N; Fig. S5B. The following antibodies were used: rabbit anti-GFP, 1:150 (Torrey Pines), guinea pig anti-Snail, 1:100 (a gift from E. Wieschaus), rat anti-DEcad2, 1:25 (Developmental Studies Hybridoma Bank, DSHB), mouse anti-Dlg, 1:50 (DSHB), rat anti-Crumbs 1:100 [Pellikka et al., 2002], mouse anti-Arm, 1:25 (DSHB), guinea pig anti-Hrs, 1:500 and guinea pig anti-Vps26, 1:500 (gifts from H. Bellen, Baylor College of Medicine, HHMI) [Wang et al., 2014], rabbit anti-beta-Galactosidase, 1:100 (Cappel), rabbit anti-Sdt, 1:3000 (gift from E. Knust, MPI CBG) [Bachmann et al., 2001], rabbit anti-PKCζ, 1:500 (C-20, Santa Cruz), guinea pig anti-Baz, 1:500 (gift from J. Zallen, MSKCC, HHMI) and rabbit anti-P-Ezrin/ERM (T567), 1:100 (Cell Signalling, 48G2). Secondary antibodies conjugated to Alexa Fluor 488, Alexa Fluor 568, or Alexa Fluor 647 (Molecular Probes) were used at 1:400.

Embryos were mounted in Prolong Gold (Molecular Probes) and imaged with zoom factors ranging from 0.75 to 3 on a TCS SP8 Leica resonant scanning confocal microscope with a HCX PL APO 63X/1.4NA CS2 objective (Leica Microsystems) via sequential scanning between channels. 1 μm Z-slices were acquired at 0.37 μm steps. Maximum projections of 2-3 μm including the apical domain or the cell nuclei were analyzed. As the rat anti-Crb antibody can detect intracellular Wolbachia (which resemble endocytic vesicles) in infected Drosophila stocks, we treated adult flies with Tetracycline at 1.5 mg/ml in food or yeast paste for 4-5 days prior to egg collection, in order to eliminate this symbiont. To select *crb*^*11a22*^ homozygous embryos expressing GFP::Crb or GFP::CrbRR (Fig. 4F), we employed LacZ staining in the progeny from *matαtub67-Gal4/+; crb*^*11a22*^ *FRT82B UAS-GFP::Crb (or UAS-GFP::CrbRR)/TM3, Sb hb-LacZ* females crossed to *crb*^*11a22*^ *FRT82B/ TM3, Sb hb-LacZ* males. Cross-section views of the ectoderm in Fig. 2E were obtained by cutting stained embryos manually with a 27-gauge syringe.

### Time-lapse imaging

Embryos expressing the indicated fluorescent markers were dechorionated for 2 minutes in 50% bleach, transferred to a drop of halocarbon oil 27 (Sigma) on a coverslip, and mounted on an oxygen-permeable membrane (YSI). GFP was excited with an OPSL 488 nm laser (2-3.5%) and mCherry was excited with an OPSL 514 nm laser (3-5%). A HCX PL APO 63X/1.4NA CS2 objective on a TCS SP8 Leica resonant scanning confocal microscope (Leica Microsystems) was used for imaging. 12-bit images of one or 2-color Z-stacks (5-13 planes; optical sections: 1.1-1.3 μm) were acquired at 0.45-0.5 μm steps every 4, 6, 10 or 15 sec intervals and maximally projected for analysis. Pixel dimensions ranged from 144 to 360 nm/pixel.

To image the consequences of the loss of Dynamin function (Dyn^TS^), *shi*^*ts1*^ embryos expressing *ubi-DEcad::GFP* or *endo-crb::GFP* were aged until mid-stage 7 (3h15 AEL) or mid-stage 8 (3h30 AEL) and imaged at 22**°**C (permissive temperature) or 32**°**C (restrictive temperature) using a stage top incubator (TOKAI HIT) assembled onto the TCS SP8 Leica resonant scanning confocal microscope.

### dsRNA and FM4-64 injections

Templates to produce dsRNA against *crb, sdt, neur and Dl* were generated by PCR from genomic DNA, using the following pairs of primers containing the T7 promoter sequence (5′-TAATACGACTCACTATAGGGAGACCAC-3′) at the 5’end:

Crb fw: 5’CGAGCCATGTCGGAATGGATCAACC 3’

Crb rv: 5’GTCGCTCTTCCGGCGGTGGCTTCAG 3’

Sdt fw: 5’ CCGTGGTACCACCGCCACTGGCGC 3’

Sdt rv: 5’ CACCCAACCCGGCCAGTTGACTGC 3’

Neur fw: 5’ CGTACGGAATCTGACTTCTGCCAGGG 3’

Neur rv: 5’ CTCGATGTACTGGCTGCTGGTGGTGC 3’

Delta fw: 5’GGAGCCTTGTGCAACGAGTGCGTTC3’

Delta rv: 5’CGCACGACAGGTGCACTGGTAATCG3’

PCR products were used as templates for T7 transcription reactions with the 5× MEGAscript T7 kit (Ambion). dsRNA was injected dorsally in 0-1 hour old embryos from the stocks *w; endo-DEcad::GFP; sqh-sqh:mCherry* (for *crb*-RNAi and *sdt*-RNAi) and progeny of *UAS-Dicer2/+*; *matαtub67-Gal4/+; matαtub15-Gal4/endo-crb::GFP* females crossed to *endo-crb::GFP* males (for *neur-*and *Dl*-RNAi).

Embryos were dechorionated, glued to a coverslip, dehydrated for 5 minutes, covered in 1:1 halocarbon oil 27/700 (Sigma), and injected with 100–200 pl of 1–2.0 μg/μl of dsRNA each. Control embryos were injected with water or not injected, as indicated. Embryos were incubated in a humidified chamber at 25°C and imaged between stages 7-9. For immunostaining, embryos were washed off the coverslip with n-heptane at stages 8 to 9, fixed for 5 minutes in 37% formaldehyde in PBS/heptane and manually devitellinized.

To visualize apical membrane internalization during ingression (Fig. 5A,B), we injected *w*^*1118*^ embryos dorsally with 100–200 pL of FM4-64 at 8 mM dissolved in 50% DMSO, either into the perivitelline space, or directly inside the embryo, and immediately imaged live using the 514 nm laser.

### Cell Segmentation and fluorescence quantification

We used SIESTA to automatically identify cell outlines in time-lapse movies using a watershed algorithm as described [Fernandez-Gonzalez and Zallen, 2015; Leung and Fernandez-Gonzalez, 2015]. When manual correction was necessary, a semi-automated method of manual tracing of cell interfaces included in SIESTA, the LiveWire, was used [Fernandez-Gonzalez and Zallen, 2013].

To measure the average apical surface area during ingression, 20-87 NBs from 2-12 embryos of each genotype were temporally aligned (registered) based on the time when they reached an apical area of 1.5 to 2.5 μm^2^. We did not consider segmentation results below this threshold range. Time 0 (onset of ingression) was defined as the time point at which the average apical surface of registered cells started declining persistently below ∼40 µm^2^, which is the average apical cell area in the ectoderm at the onset of ingression during stage 8 [Fernandez-Gonzalez and Zallen, 2015]. To compare rates of ingression, we matched initial average areas in control and mutant/RNAi embryos. Ingression speed of individual NBs was the slope of a linear fit (first degree polynomial) for the apical area loss over time (using the Matlab function *polyfit*). When average fluorescence results from multiple cells were analyzed, cells were temporally registered using the same area threshold. For all experiments, we compared controls to mutant or RNAi embryos carrying the same fluorescent marker(s), and imaged with the same settings and environmental conditions.

To quantify junctional and medial average protein levels, each cell was divided into two compartments as described [Fernandez-Gonzalez and Zallen, 2015]. The junctional compartment was determined by a 3-pixel-wide (0.54 μm) dilation of the cell outline identified using watershed or LiveWire segmentation. The medial compartment was obtained by inverting a binary image representing the junctional compartment. Protein concentrations were quantified as the mean pixel intensity in each compartment considering either all ingression time points, up to the last 10 minutes of the process (early ingression), or the last 10 minutes (late ingression), as indicated.

When determining total or average protein intensities, we combined results from different embryos of the same genetic background/fluorescent marker, imaged with the same confocal settings, and at the same temperature. Average fluorescence intensities were normalized at each time point by subtracting the fluorescence mode for the entire image (background) and dividing by the mean pixel value of each frame. When comparing different genetic backgrounds expressing the same fluorescent marker (control *versus* mutant or RNAi), average fluorescence intensities at each time point were normalized by subtracting the fluorescence mode for the entire image (background) at that time point, on all movie time frames.

Total Crb and Ecad intensity at each time point were the sum of pixel intensities in the junctional domain normalized as indicated above. Relative changes in apical cell perimeter and total apical Crb levels throughout ingression (Fig. 1C and D) were defined as:

Δperimeter (t)= (Apical Perimeter (t) – Apical Perimeter (t-60 sec))/ Apical Perimeter (t-60 sec) Δtotal Crb levels (t)= (Total protein (t) – Total protein (t-60 sec))/ Total protein (t-60 sec)

To determine fractions of Crb loss or gain during apical oscillations (Figs. 1F, S1A), the apical perimeter and total fluorescent levels of Crb in individual NBs were smoothened by averaging over 2 consecutive data points at a temporal resolution of 15 sec. “Peaks” and “troughs” were then identified from smoothened perimeter values by imposing a minimum separation of 30 seconds between consecutive peaks and troughs and a perimeter change of at least 1 μm. Contractions were defined as apical oscillations from consecutive “peaks” to “troughs” and expansions were defined as apical oscillations from consecutive “troughs” to “peaks”. We calculated fractions of perimeter or Crb change during contractions and expansions as:

Fraction_perimeter_change_contraction = (Perimeter(peak) – Perimeter(trough))/ Perimeter(peak)

Fraction_total_Crb_change_contraction = (Total_Crb(peak) – Total_Crb(trough))/ Total_Crb(peak)

Fraction_perimeter_change_expansion = (Perimeter(trough) – Perimeter(peak))/ Perimeter(trough)

Fraction_total_Crb_change_expansion = (Total_Crb(trough) – Total_Crb(peak))/ Total_Crb(trough)

Rate of apical area change, ΔArea (t), was defined as ΔArea (t)= Area (t) – Area (t-60 seconds), whereas rate of medial or junctional myosin II change was defined as Δmyosin (t)= myosin concentration(t) – myosin concentration (t-60 seconds). The duration of apical contractions and expansions was defined as the number of consecutive time points multiplied by time resolution during which ΔArea(t) was <0 or >0, respectively. The amplitude of contractions and expansions was the maximum value of area change within each event.

To quantify FM4-64 internalization in live NBs and NICs, we quantified the average FM4-64 fluorescence in the medial cell compartment throughout time after injection (images acquired at 4 seconds intervals). The image intensity mode (background) was subtracted at each time point, and the result divided by the image intensity mean at each time point. Fluorescence intensities were then normalized to the first time point of movie acquisition, in order to compare the relative internalization of FM4-64 in ingressing NBs versus NICs.

To determine total levels of internalized Crb in fixed NBs, we employed LiveWire in Siesta and segmented the membrane outline of NBs at individual Z planes (Z step size = 0.37 μm) encompassing the entire apical-basal height of the cell. Total intracellular Crb levels were the sum of total fluorescence intensity in the medial compartment for all Z planes encompassing that cell. Fluorescence intensities were normalized by subtracting the fluorescence mode at each Z plane. The size of Crb containing endosomes in NBs and non-ingressing cells was determined in Fiji using Watershed Segmentation and Analyze Particles in manually drawn ROI around cells of interest. Original images in the Crb channel were converted to binary images via thresholding (default method in Fiji); only segmented vesicles with a larger size than 10 pixels (1 pixel=60.13 nm) were considered for analysis. Data was obtained from wild-type embryos using fixation Method B (see above).

For colocalization analysis between internalized Crb and several endocytic/cytoskeletal/polarity markers (Figs. 4B,D and S3), we used Coloc2 (Fiji) in manually drawn ROIs around NBs of fixed embryos, in order to obtain Pearson’s coefficients and Mander’s coefficients. We employed the Costes method [Costes et al., 2004] to assess the significance of the calculated Mander’s co-localization coefficients, by analysing 200 image randomizations in the Crb channel for each cell, considering a point spread function (PSF) of 3 pixels (1 pixel = 60.13 nm).

### Statistics

Average or median values were determined based on a certain number, n, of embryos/cells/ /contractions/expansions, as indicated in each figure legend. Error bars are SD (standard deviation) or SEM (standard error of the mean), which is SD/ √n or interquartile ranges, as indicated. We used Graphpad Prism 8 to test if the n values of each sample followed a normal distribution, by performing a D’Agostino and Pearson normality test. We employed unpaired 2-tailed T tests to determine p values when samples passed the normality test and non-parametric 2 sample Kolmogorov Smirnov (KS) or Mann-Whitney tests otherwise. F-test was used to assess the variance between two samples.

## Acknowledgements

We thanks Francois Schweisguth, Hugo Bellen, Yang Hong, Chris Doe, Elisabeth Knust, Adam Martin, Eric Wieschaus, Jennifer Zallen, Hiroki Oda, Thomas Lecuit, Daniel St. Johnston, the Bloomington Drosophila Stock Center, the Drosophila RNAi Screening Center at Harvard Medical School, and the Developmental Studies Hybridoma Bank for reagents. We like to thank Dorothea Godt for critical reading of the manuscript, and the Imaging Facility of the Department of Cell and Systems Biology, University of Toronto, for support. **Funding:** This work was funded by an Innovation Grant from the Canadian Cancer Society (to U.T. and R.F.G.) and a project grant from the Canadian Institutes for Health Research (to U.T. and R.F.G.). R.F.G. is a Canada Research Chair in Quantitative Cell Biology and Morphogenesis and U.T. is a Canada Research Chair for Epithelial Polarity and Development.

## Author contributions

The project was conceived and experiments were designed by S.S. and U.T. S.S. did the majority of the experimental work and data analysis. G.L., M.P., D.t.S, and K.A.K. contributed to the experimental work. G.L. P.G., T.L., and D.K. contributed to data analysis. J.Y. and R.F.G. generated code and contributed expertise in data analysis. R.F.G and U.T. provided supervision and raised funds. The paper was written by S.S. and U.T.

## Data and material availability

All data are available in the manuscript or supplementary materials.

## Supplementary materials

Materials and Methods

Supplemental Figures S1 – S5

Supplemental Videos 1 – 10

