## Supplementary materials for "Crumbs complex-directed apical membrane dynamics in epithelial cell ingression"

February 23, 2022

### Supplementary Materials

- Supplemental figure legends (S1-S5)
- Supplemental Video legends (1-10)

**Figure S1. Crb dynamics during apical contractions, and NB apical area, myosin, and Ecad dynamics in *crb*-RNAi embryos.**

(**A**) Left dot plot showing the fraction of apical perimeter increase (exp, expansions) or decrease (cont, contractions) during early and late ingression. ns, p=0.14; ****, p=3X$10^{-9}$, 3.17X$10^{-6}$ and 4.6X$10^{-12}$. Right dot plot showing the fraction of apical Crb gain or loss during apical oscillations in early and late ingression. ns, p=0.78 and 0.19; ****, p=8.57 X$10^{-10}$; ***, p= 3X$10^{-4}$. Note a significant increase in relative contractions during late ingression, matching with higher reductions in apical Crb. Column height, median; error bars, IQR; 385 contractions and 302 expansions analyzed in 58 NBs from 12 embryos expressing endo-Crb::GFP.

**(B)** Ventral views of late stage 8 embryos injected with $H_2O$ (top, 4 embryos) or *crb*-dsRNA (bottom, 7 embryos) stained for Crb and Armadillo (Arm). Notice the strong uniform reduction in Crb levels in *crb*-RNAi embryos. Scale bars, 50 μm.

**(C)** Stills from time-lapse movies of ingressing NBs (arrowheads) expressing Ecad (endo-Ecad::GFP, green) and myosin::mCherry (sqh-Sqh::mCherry, red) in *crb*-RNAi and embryos ($H_2O$ injected control is shown in Figure 2A). T=0 min, onset of ingression. Scale bars, 5 μm.

**(D)** Amplitude and duration of individual contractions/expansions during ingression in control versus *crb*-RNAi slowest and fastest NBs. Median amplitudes for control/slow/fast *crb*-RNAi of contractions, 3.04/2.35/6.95 $\mu m^2$/min; **, p=7.6×$10^{-3}$; ****, p=1.5X$10^{-16}$ and 2.0X$10^{-17}$; and expansions, 1.42/1.32/2.4 $\mu m^2$/min; ns (not significant), p = 0.23; ****, p=3.3X$10^{-5}$ and 5.4X$10^{-6}$. Median duration for control/slow/fast *crb*-RNAi of contractions, 46.6/38.3/53.2 sec, ns, p=0.10; *, p = 0.046; **, p=5.4X10-3; and expansions, 20.6/21.2/19.6 sec, ns, p = 0.09, 0.16, 0.06 (KS test); n = 176–551 events per condition.

**(E)** Representative plots showing the rate of medial myosin change (a.u./min) throughout ingression in a control NB (blue), and *crb*-RNAi slow and fast NB (red). Positive rates indicate myosin assembly; negative rates indicate disassembly. T=0 min, onset of ingression.

**(F)** Medial myosin assembly and disassembly rates during ingression in control NBs, *crb*-RNAi slow and fast NBs. N values as in Fig. 3C. Median assembly rates for control/slow/fast Crb RNAi: 0.06/0.09/0.09

a.u./min; ***, $p=5.0\times10^{-4}$; *, p=0.01; median disassembly rates for control/slow/fast Crb RNAi: 0.05/0.09/0.06 a.u./min; ***, $p=1X10^{-4}$; ns (not significant), p = 0.07; n=151-429 events per condition.

**(G)** Mean levels of Ecad (endo-Ecad::GFP) in control NBs and slow/fast *crb*-RNAi NBs. N values as in Fig. 3C. ****, $p=1.8X10^{-10}$, $8.3X10^{-79}$, $1.7X10^{-49}$ (2-tailed T test). Means plus SD.

**Figure S2. Junctional and medial myosin dynamics in *crb*-RNAi NBs versus NICs.**

**(A, B)** Junctional (A) and medial (B) myosin ratios between ingressing NBs and their surrounding non-ingressing cells (NICs) (schematics), plotted against ingression speed (apical area loss in $\mu m^2$/min). Dots are the medians of myosin ratios NB/$NIC_{A-F}$, determined while the NB apical surface decreased from 20 $\mu m^2$ to 2.5 $\mu m^2$. Bars are IQRs. N=19 control NBs, 3 embryos; N=25 *crb*-RNAi NBs, 5 embryos. $R^2$ is the correlation coefficient of a linear fit, with its corresponding p value. ns, not significant.

**Figure S3. Colocalization analysis of endocytic Crb in ingressed NBs.**

**(A-N)** Single ingressed NBs co-stained for Crb, Dlg (lateral membrane marker) and the indicated markers in green. (A) and (B) are positive and negative co-localization controls, respectively: NBs from endo-Crb::GFP expressing embryo were co-stained for Crb and GFP. The GFP channel was rotated 90° clockwise in (B). Scale bars, 2.5 μm.

**(O)** Mander's colocalization coefficients (orange bars, mean +/- SD) between intracellular Crb and markers indicated in (A-N) and in Fig. 3b. Blue bars indicate the fraction of NBs with PCostes >0.95. Mander's coefficients were determined when more than 80% of NBs had PCostes >0.95 (markers to the left of dashed line). +/- control: 90 NBs, 7 embryos; Rab5: 128 NBs, 7 embryos; P(I)3P: 238 NBs, 6 embryos; Rab7: 159 NBs, 6 embryos; Sdt: 177 NBs, 7 embryos; Lamp-1: 119 NBs, 4 embryos; Rab11: 89 NBs, 4 embryos; Par6: 99 NBs, 4 embryos; aPKC: 119 NBs, 6 embryos; Par3: 145 NBs, 6 embryos; PIP2: 42 NBs, 4 embryos; β Heavy-Spectrin: 101 NBs, 5 embryos; Phospho-Moesin: 95 NBs, 4 embryos.

**(P)** Pearson's colocalization coefficients between intracellular Crb and the indicated markers in ingressed NBs. Bars indicate mean +/- SD. N values as in (O).

**(Q,R)** Representative scatter plots of individual embryos displaying the relationship between apical surface and intracellular Crb levels in ingressing NBs (blue dots). (Q) Late stage 8 embryo (ingression is underway), 33 NBs. (R) stage 9 embryo (ingression is complete in most NBs), 16 NBs. Note an increase in intracellular Crb as NBs reduce their apical surface. Fully ingressed cells (with apical surface ~ 0 $\mu m^2$) may display variable levels of intracellular Crb likely reflecting post-ingression Crb protein degradation. Purple lines represent the cubit fit of experimental data.

**Figure S4. Dynamin is required for the loss of the apical domain during NB ingression.**

**(A)** Stills from time-lapse movies of ingressing NBs of $Dyn^{TS}$ ($shi^{ts1}$ mutants) at 22°C and 32°C, expressing ubi-Ecad::GFP. Dividing ventral midline cells (VMC) are pseudo-colored in blue. Note ingression of medial NBs (yellow circles) adjacent to the VMC at the permissive temperature of 22°C. Ingression is stalled upon Dynamin inhibition at the restrictive temperature of 32°C and ectopic Ecad accumulation is seen at cell-cell contacts throughout the germband No NBs with constricted apical domains are apparent. T=0 min indicates onset of stage 8. Scale bar, 10 μm.

**(B)** Stills from time-lapse movies of ingressing NBs of $Dyn^{TS}$ at 22°C (permissive temperature; 32 NBs, 3 embryos) and 32°C (restrictive temperature; 41 NBs, 2 embryos) expressing endo-Crb::GFP. Dynamin blockage at the restrictive temperature beginning at mid-stage 8 (3h30min after egg laying) led to incomplete ingression in 56% of NBs and 22% of NBs divided on the embryo surface (normal divisions occur after NB complete ingression and are located below the ectoderm). Note higher accumulation of Crb on the apical surface of the delayed cells at 32°C compared to controls at 22°C (quantifications shown in Fig. 4C). Scale bar, 2.5 μm.

**Figure S5. Loss of Neur causes apical Crb accumulation and slows NB ingression.**

(**A**) Live stills from un-injected embryos (control) or *sdt* dsRNA injected embryos (*sdt*-RNAi), expressing endo-Sdt::GFP (5 embryos each), GFP::Crb (n=5 and 8 embryos, respectively) and GFP::CrbRR (n=8 and 6 embryos, respectively). In the absence of Sdt, GFP::CrbRR remains at the plasma membrane in contrast to GFP::Crb, which is found in endosomes. Scale bar, 10 μm.

**(B)** Ventral views of late stage 8, early stage 9 and mid stage 9 embryos expressing Dicer-2 maternally and injected with water (control, n=13) or with *neur* dsRNA (*neur*-RNAi, n=12), stained for Crb and Moesin. Brackets delimit the neuroectoderm. Note ingression of NBs in clusters enriched for apical Crb in *neur*-RNAi embryos (insets), whereas in controls NBs ingress as individual cells. Ingression ends by early stage 9 (inset, control), whereas apical NB clusters are still visible in *neur*-RNAi embryos at that stage. Upon division, these supra-numerary NBs disrupt neurectoderm integrity. Anterior is to the left, dorsal is up. Scale bar, 50 μm.

**Supplementary Videos:**

1. *matαtub67-Gal4; matαtub15-Gal4, UAS-GFP::crb* (control)
2. *matαtub67-Gal4; matαtub15-Gal4, UAS-GFP::crbRR* (NB, surface division)
3. *matαtub67-Gal4; matαtub15-Gal4*, *UAS-GFP::crbRR* (NB, slow ingression)
4. Injection of FM4-6, 16mM
5. Injection of FM4-6, 8mM
6. *matαtub15-Gal4*, *GAP43::mCherry* (control)
7. *matαtub15-Gal4*, GAP43::mCherry, *UAS-AP2α RNAi* (slow ingression)
8. *matαtub15-Gal4*, GAP43::mCherry, *UAS-AP2α RNAi* (surface division)
9. *endo-sdt3::GFP*
10. *endo-sdtΔ3::GFP*

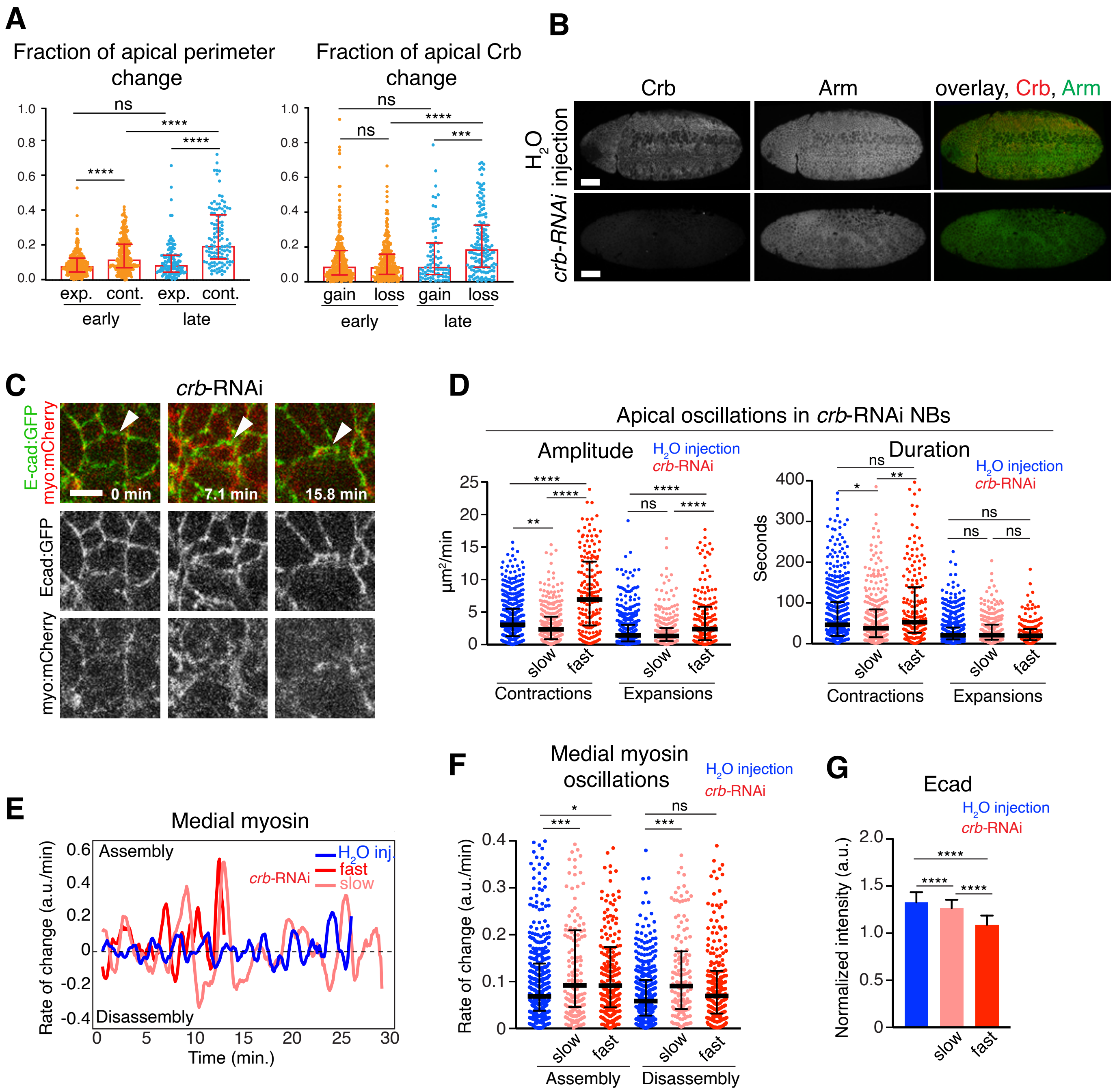


**Simoes et al. Fig. S1**

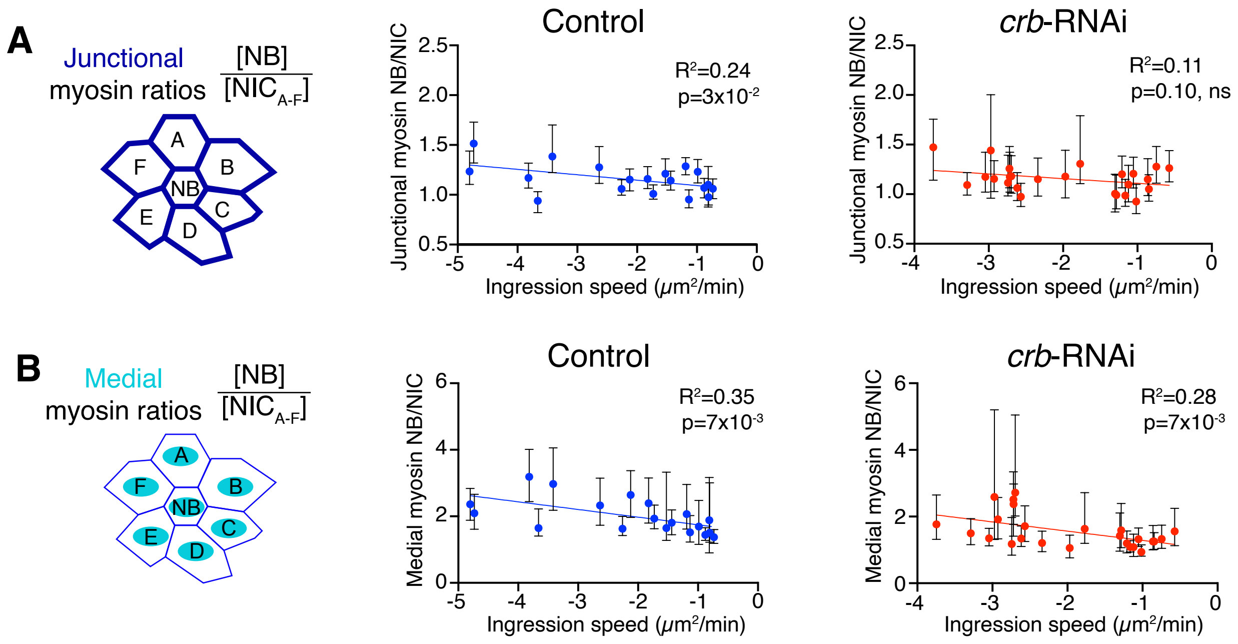


**Simoes et al. Fig. S2**

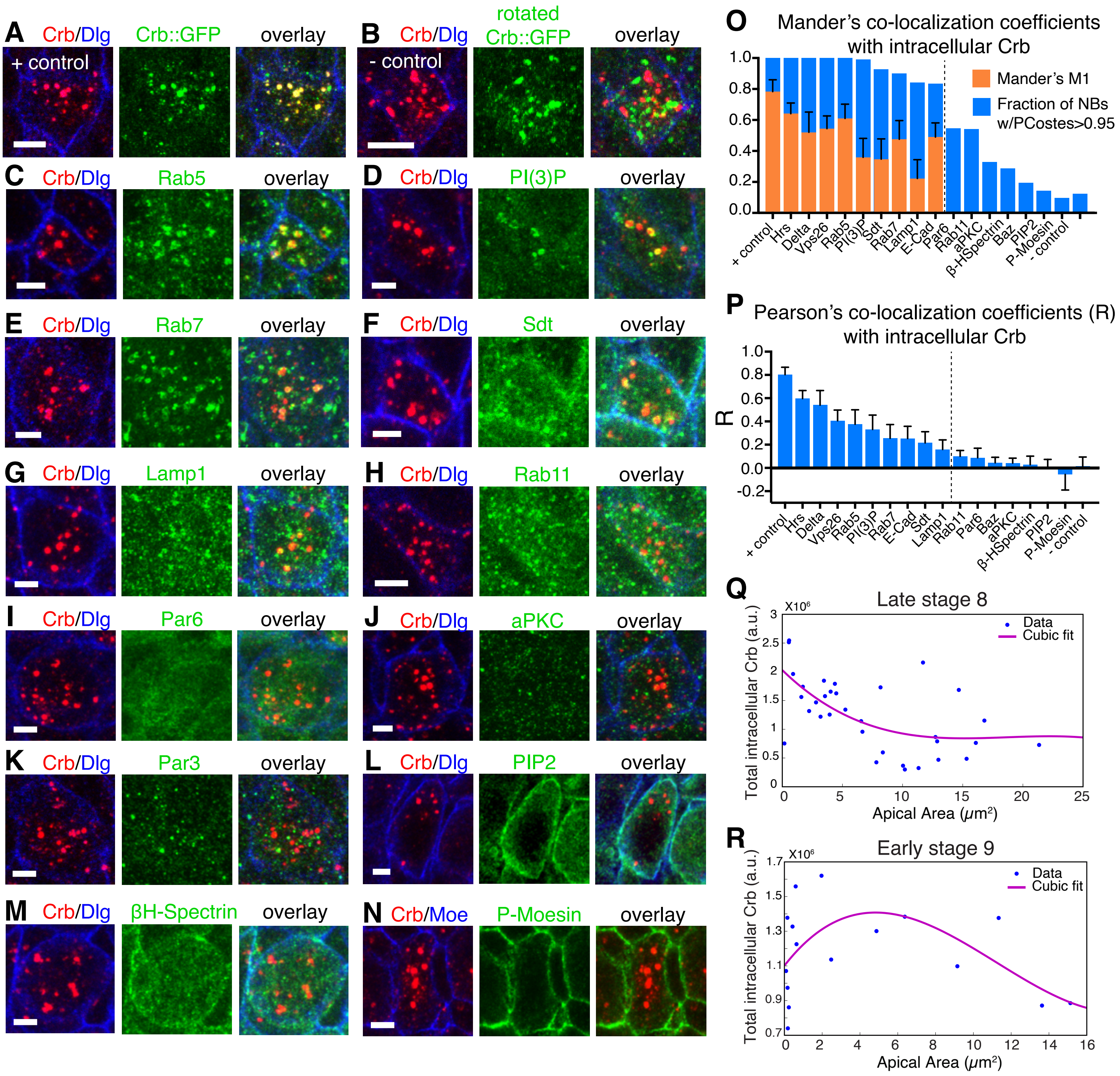


**Simoes et al., Fig. S3**

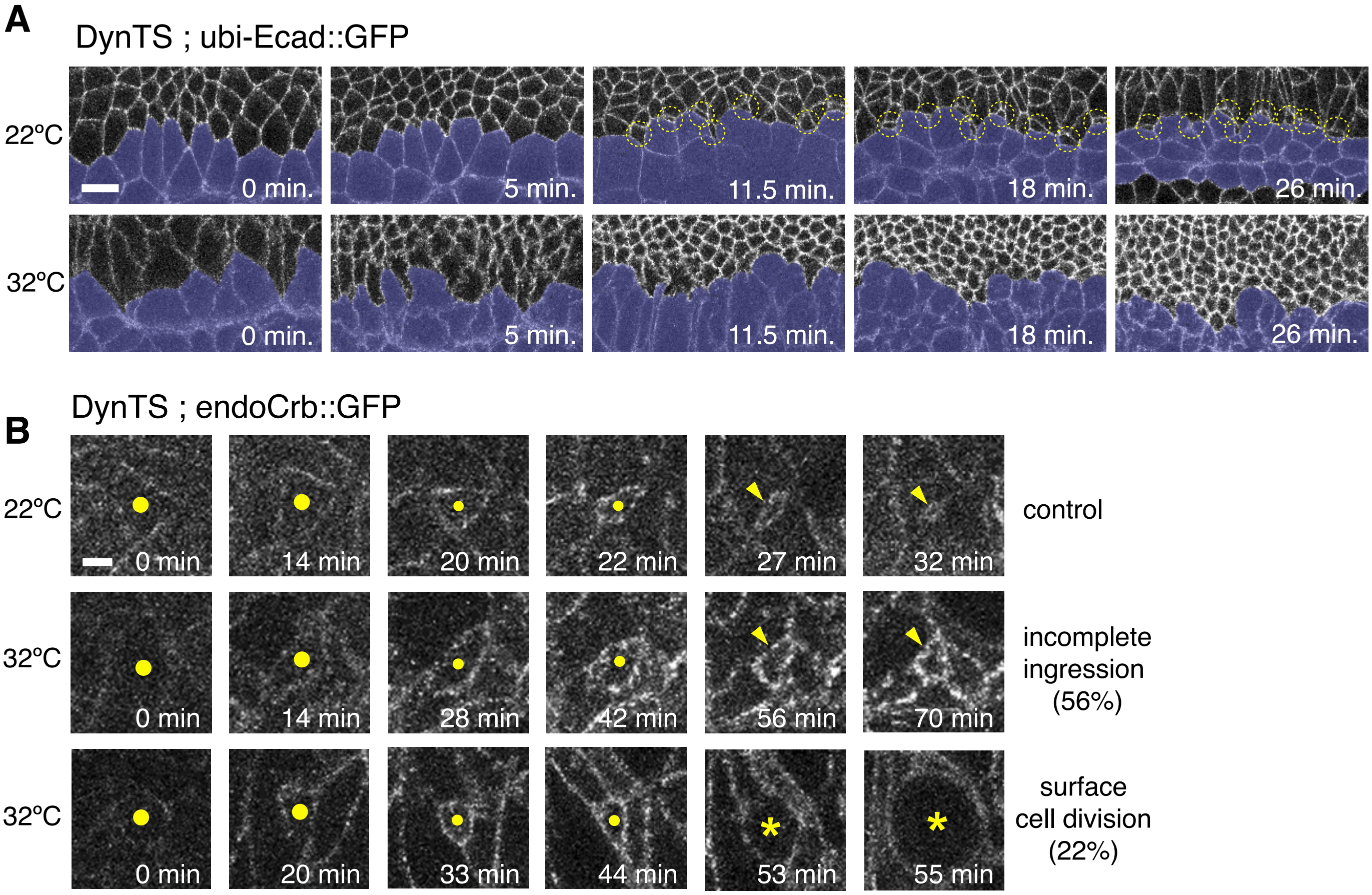


**Simoes et al. Fig. S4**

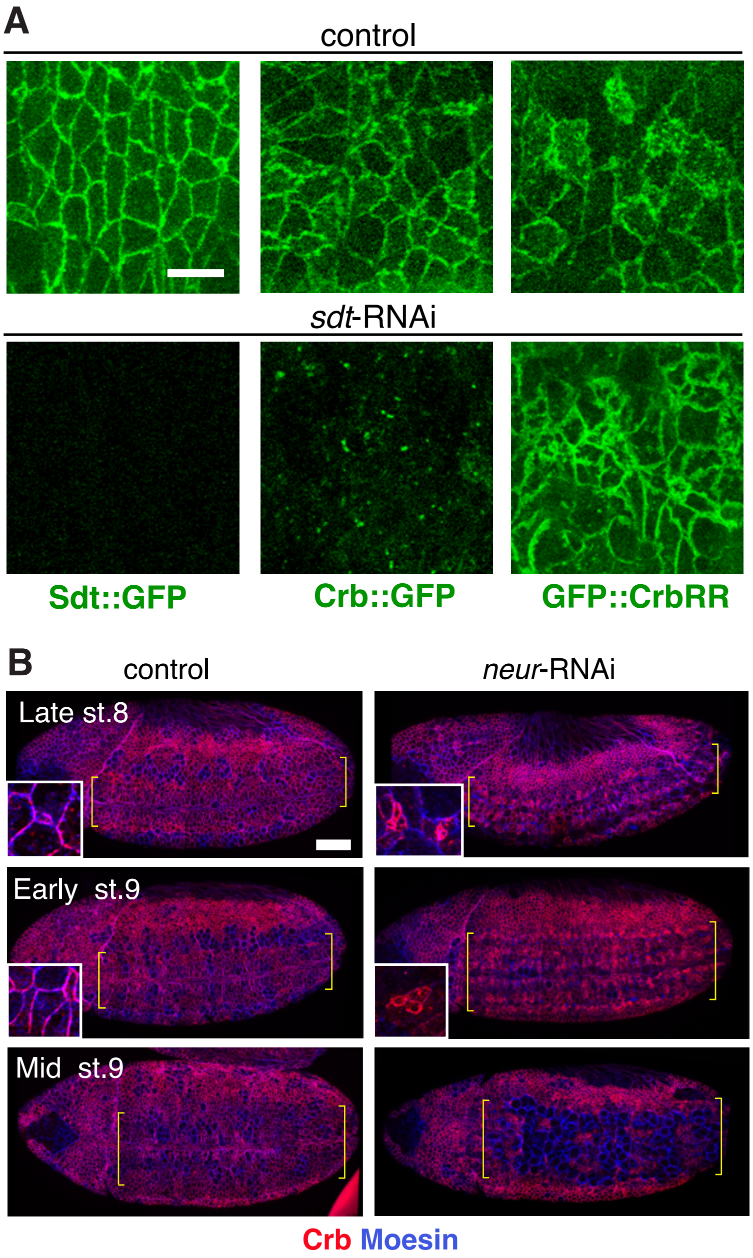


**Simoes et al. Fig. S5**
